# A comprehensive benchmark for multiple highly efficient base editors with broad targeting scope

**DOI:** 10.1101/2024.12.17.628899

**Authors:** Xiaofeng Wang, Xiaolong Cheng, Zexu Li, Shixin Ma, Han Zhang, Zhisong Chen, Yingjia Yao, Zihan Li, Chunge Zhong, You Li, Yunhan Zhang, Vipin Menon, Lumen Chao, Wei Li, Teng Fei

## Abstract

As the toolbox of base editors (BEs) expands, selecting appropriate BE and guide RNA (gRNA) to achieve optimal editing efficiency and outcome for a given target becomes challenging. Here, we construct a set of 10 adenine and cytosine BEs with high activity and broad targeting scope, and comprehensively evaluate their editing profiles and properties head-to-head with 34,040 BE-gRNA-target combinations using genomically integrated long targets and tiling gRNA strategies. Interestingly, we observe widespread non-canonical protospacer adjacent motifs (PAMs) for these BEs. Using this large-scale benchmark data, we build a deep learning model, named BEEP (<u>B</u>ase <u>E</u>diting <u>E</u>fficiency <u>P</u>redictor), for predicting the editing efficiency and outcome of these BEs. Guided by BEEP, we experimentally test and validate the installment of 3,558 disease-associated single nucleotide variants (SNVs) via BEs, including 20.1% of target sites that would be generally considered as “uneditable”, due to the lack of canonical PAMs. We further predict candidate BE-gRNA-target combinations for modeling 1,752,651 ClinVar SNVs. We also identify several cancer-associated SNVs that drive the resistance to BRAF inhibitors in melanoma. These efforts benchmark the performance and illuminate the capabilities of multiple highly useful BEs for interrogating functional SNVs. A practical webserver (http://beep.weililab.org/) is freely accessible to guide the selection of optimal BEs and gRNAs for a given target.

## Introduction

The advent of base editors (BEs) enables precise conversion of target nucleotides in the genome without requiring double-strand breaks and donor DNA templates^1, 2^. Given that single-nucleotide variants (SNVs) represent the major types of genomic alternations in many genetic disorders and cancer^3, 4^, BE provides a simple yet effective tool in both basic research and therapeutic applications to install or correct SNVs. Classic BEs are usually composed of a catalytically impaired Cas protein and a base-modifying enzyme. The Cas9 domain directs BE to the target site, and the deaminase domain catalyzes the conversion of targeted nucleotide bases. Adenine base editor (ABE) converts A•T-to-G•C, whereas cytosine base editor (CBE) converts C•G-to-T•A^1, 2^. Recently, other types of BEs were also developed to mediate more diverse types of nucleotide conversion (e.g., C•G-to-G•C, A•T-to-T•A or A•T-to-C•G)^5–13^ or constructed under Cas protein-independent guiding systems such as IscB-ωRNA^14–19^. Rational design, directed evolution, and artificial intelligence (AI) further contribute to the rapid expansion of the BE toolbox with varied editing efficiency, targeting scope, editing profile, and other properties^20–38^. However, the large number of available BEs, without comprehensive and rigorous benchmarking, makes it challenging for the end-users from different fields to select appropriate BE and suitable guide RNA (gRNA) for their base editing needs.

Traditionally, BEs are usually evaluated using a limited number of target sites and/or an array of target-gRNA catenated fragments in a high-throughput manner^39–47^. The target fragments are generally short (∼ tens of nucleotides), and the variety by design mainly lies in the subregions around protospacer adjacent motif (PAM) or adjacent to target nucleotides. Aided by AI and deep learning models, such strategy has proven to be effective in learning global features that determine the editing outcomes for a given BE, such as PAM preference, editing context, mismatch tolerance, editing window, bystander editing, product purity, etc.^39, 41, 46, 47^. However, it remains challenging to directly compare the performances of multiple BEs head-to-head, because that 1) different BEs are often characterized with different custom-designed target-gRNA pairs; 2) the expression levels of different BEs are usually not strictly calibrated; and 3) short target-gRNA fragment still lacks sufficient context information, especially when unbiasedly comparing the editing outcomes of different BEs in editing the same target region.

Here, we construct a set of 10 adenine and cytosine BEs with high activity and broad targeting scope which can efficiently cover as many target sites as possible for ABE/CBE. Using the same genomically integrated long targets and tiling gRNA libraries in BE expression-controlled human cells, we benchmark these BEs head-to-head to evaluate their editing profiles and properties. Using these BEs, we prove their collective power to successfully install or interrogate pathological or cancer-associated SNVs. We further build a deep learning model BEEP and a practical webserver to assist end-users in selecting optimal BE-gRNA combination for a given target.

## Results

### Long targets and gRNA tiling strategy for benchmarking a set of BEs with high usability

To develop a BE toolbox with maximal targeting scope at high editing efficiency, we constructed a set of BEs by combining Cas9 variants of differential PAM preferences (SpCas9 - NGG PAM; xCas9 - NG, GAA, and GAT PAM^48^; Cas9-NG - NG PAM^49^; SpG - NGN PAM^49^; SpRY – NRN > NYN PAM^50^) with two of the most effective deaminase effectors to date (ABE8e and evoAPOBEC1-BE4max)^23, 27^. Human codon-optimized sequences of the above domains were cloned into a lentiviral vector with indicated architectures (Fig. 1a and Extended Data Fig. 1a; Supplementary Note 1), giving rise to the following BEs: five ABEs (SpCas9-ABE8e, xCas9-ABE8e, Cas9-NG-ABE8e, SpG-ABE8e, and SpRY-ABE8e) and five CBEs (SpCas9-evoA-BE4max, xCas9-evoA-BE4max, Cas9-NG-evoA-BE4max, SpG-evoA-BE4max, and SpRY-evoA-BE4max). Using a low multiplicity of infection (MOI) (∼0.1) to transduce these BEs into human melanoma A375 cells at equally low copies via lentiviral infection, we were able to compare their expression levels in mammalian cells between different BEs. Despite variations, all the BEs have comparable levels in cells even at a low MOI (Extended Data Fig. 1b). We further transduced gRNA-expressing lentiviral particles still at a low MOI (∼0.3) into these BE-expressing A375 cells to examine their base editing capabilities on individually selected genomic sites. As expected, for target sites containing a classic NGG PAM, all these 10 BEs showed strong yet variable base conversions (∼20%-60% edit frequency) within target windows between 2 to 11 (counting the position immediately upstream of the protospacer as position 1) (Extended Data Fig. 1c). However, for those target sites without classic NGG PAMs, the editing efficiencies varied a lot across different BEs. Some Cas9 variant-derived BEs outperformed the classical SpCas9-based ABE or CBE in a loci- and sequence-dependent manner (Extended Data Fig. 1c), highlighting the need to broaden the targeting scope and predict the editing outcome with new BEs.

**Fig. 1.**
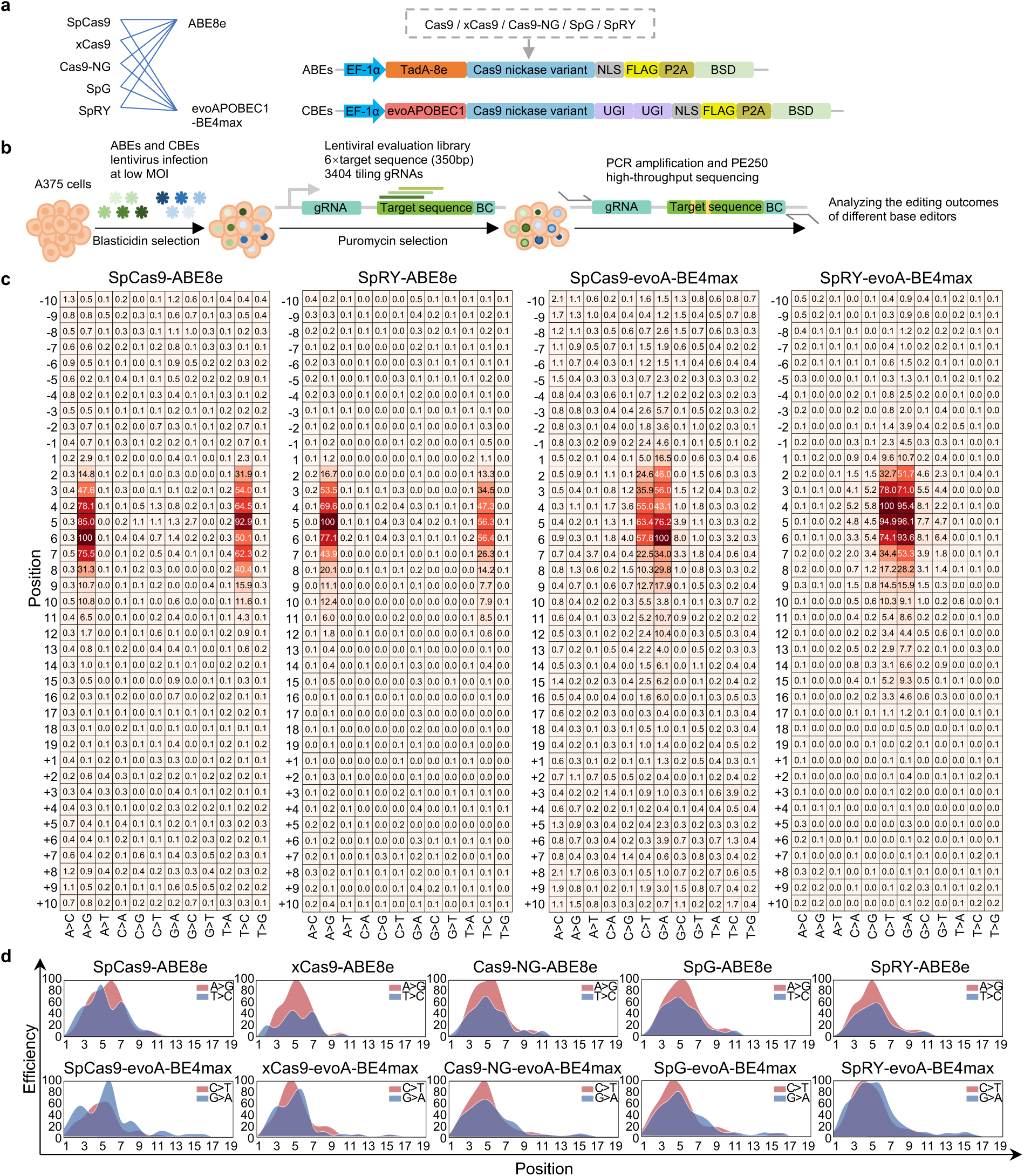
High-throughput evaluation of base editing outcomes via long targets and gRNA tiling strategy. **a**, Schematic representation of base editors constructed via combination of different deaminase effectors and Cas9 variants. **b**, Schematic representation of the strategy for high-throughput evaluation of these ABEs and CBEs. A375 cells were firstly infected with lentiviruses at low MOI to generate expression-calibrated ABE- or CBE-expressing cell lines. The subsequent delivery of lentiviral BE evaluation library containing matched gRNA-target-barcode sequences into these cells induced base conversion at the genomically integrated target sequences. The whole fragment was PCR amplified for PE250 high-throughput sequencing to measure the editing outcomes. **c**, Heatmap showing the base editing window for indicated BEs. A>G/C>T indicates the spacer (gRNA) and target aligning at the same strand, while T>C/G>A means the spacer and target aligning at different strands. Values in the heatmap indicate the normalized editing efficiency (percentage to the maximal editing). **d**, The target position and editing efficiency relationship for default A>G/C>T conversion on the same strand or complementary target nucleotide T>C/G>A conversion on the different strands.

To comprehensively compare the performance of different BEs, we synthesized an oligo library with 3,403 tiling gRNAs targeting the whole EGFP and luciferase coding sequences regardless of any PAM restriction (Supplementary Data 1). Using a two-step cloning method, we incorporated these oligos into a lentivector backbone to produce gRNA-expressing lentiviral library along with their cognate target fragments (Extended Data Fig. 1d; see Methods). Each gRNA is concatenated with a 350-base pair (bp) EGFP or luciferase fragment containing its cognate target region, followed by a unique 15 bp barcode sequence (Fig. 1b). We separately transduced this library into the ten A375 lines expressing each one of the BE systems at low MOI (∼0.3) followed by puromycin selection. The cells were continuously cultured for 7 days to allow for genomic integration of gRNA-target fragment cassette and base conversion, and high-throughput long-read paired-end 250 (PE250) sequencing was deployed to determine the abundance and sequence identity of the whole gRNA-target-barcode region (see Methods). Unlike previous studies, our experimental design, with the same unbiased tiling gRNA library and no PAM restriction, long-read sequencing mode, and expression-controlled BE-expressing cells, enabled us to comprehensively and unbiasedly benchmark the performances (e.g., editing efficiency, PAM preference, sequence context, etc.) of different BEs. A total of 20 high-throughput evaluation assays were performed with two biological replicates for each BE group. After removing the reads derived from sequencing errors and multi-stage recombination or uncoupling (see Methods), the editing frequency and outcome of target sites in the evaluation library were analyzed. The two biological replicates were highly correlated (average Pearson’s *r* = 0.90 across all the conditions) in terms of the editing frequencies of targets with cognate PAMs for each BE (Extended Data Fig. 1e), suggesting a high reproducibility and good quality of these data.

### Editing profiles and properties of various BEs

Using the data from the above-mentioned high throughput evaluation assays, we quantified the base editing outcomes of all possible nucleotide substitutions spanning - 10 to +10 of protospacer positions for all the 10 BEs to obtain their global editing profiles (Fig. 1c,d and Extended Data Fig. 2a,b). Although both ABEs and CBEs displayed highest editing activities toward corresponding A•T-to-G•C and C•G-to-T•A conversions, respectively, different BEs still varied in strongest editing position, editing window width and non-canonical editing patterns. For instance, classic SpCas9-ABE8e showed highest A•T-to-G•C conversion at position 6 (Fig. 1c), whereas other Cas9 variant-derived ABEs all displayed peak editing at position 5 (Fig. 1c and Extended Data Fig. 2a). For all the BEs evaluated, the prominent activity window was positioned between position 3 to 7; however, CBEs tended to extend their editing territories even outside of protospacers to upstream position −1 or −2 (with >2% relative editing frequency) while ABEs did not (Fig. 1c,d and Extended Data Fig. 2a,b). In addition to standard base conversion types (A•T-to-G•C for ABEs; C•G-to-T•A for CBEs), CBEs generally possessed more non-canonical editing patterns (considering >1% relative editing frequency as an arbitrary cutoff) and higher strength than ABEs. Except for SpCas9-evoA-BE4max, all the other four CBEs exhibited strong non-canonical C•G-to-A•T or C•G-to-G•C conversion (>5% relative editing frequency). In contrast, only SpCas9-ABE8e and xCas9-ABE8e displayed a modest non-canonical G•C-to-A•T or G•C-to-C•G editing, the other three ABEs barely had any non-canonical base conversion (Fig. 1c and Extended Data Fig. 2a,b). These results suggest that BE-conjugated Cas9 variants not only determine the targeting scope (PAM selection), but also affect the catalytic processes of deaminases and the resulting base conversion patterns. Interestingly, whether the target nucleotide in gRNA aligns in the same strand direction as the target nucleotide in genomic DNA affects the editing efficiency and activity window for all the BEs. In other words, when editing the complementary nucleotide (T>C for ABE; G>A for CBE) on the default strand, the editing rules would be different from those gained from the target nucleotide in the default strand (A>G for ABE; C>T for CBE) (Extended Data Fig. 2c,d). For example, when the spacer (gRNA) and target genome were in the same strand direction for SpRY-ABE8e, the target nucleotide ‘A’ in the default strand will be converted to ‘G’ with the highest relative efficiency at position 5. On the other hand, T-to-C transition in the default strand can be achieved by A-to-G editing on the opposite strand with the same SpRY-ABE8e (Extended Data Fig. 2c,d). However, in such T•A-to-C•G (T>C) transition scenario, the editing efficiency at peak position dropped significantly compared to A•T-to-G•C (A>G) conversion (56.4% at position 6 vs. 100% at position 5) (Fig. 1c). Another interesting case was SpCas9-evoA-BE4max, which rather displayed stronger editing efficiency for G•C-to-A•T (G>A) transition compared to the default C•G-to-T•A (C>T) conversion (Fig. 1c). Such discordance of editing properties between target and complementary nucleotides on the same default strand may be due to different reasons, including 1) different steps and mechanisms (such as DNA repair and/or replication) are required to complete target or complementary nucleotide conversion (Extended Data Fig. 2c,d), and 2) undefined factors during these processes might affect the efficiencies of different base substitution. This finding is important as new rules should be considered when using these BEs to edit complementary nucleotide T or G on default strand by ABE or CBE in many scenarios.

Next, we extensively examined different aspects of editing properties for all the ten BEs. As the expression levels of BE and gRNA were carefully gauged by introducing the same minimal copies of lentiviral particles with a low MOI, we were able to directly compare the editing efficiencies between different BEs. For most ABEs, the absolute A•T-to-G•C (A>G) transition efficiencies (counting both default strand and complementary strand editing together) at peak positions reached 30%∼40%, except for xCas9-ABE8e with relatively lower peak activity (20%∼30%) (Fig. 2a). All the CBEs generally showed a slightly weaker peak editing efficiencies (20%∼30%) in converting C•G-to-T•A (C>T) than those of A>G transition by ABEs. Among all CBEs, xCas9-evoA-BE4max showed weakest efficiency (Fig. 2a). On the other hand, the active editing window of CBEs was generally wider than that of ABEs (Fig. 2a). To learn the sequence context that affects the editing efficiencies (A>G for ABE; C>T for CBE), we calculated the sequence motifs that are enriched in guides (gRNA matched protospacer sequences) with higher editing efficiencies. All the ABEs shared a similar upstream GC<u>A</u> motif but slightly differed at downstream motifs, especially at position +2 (G or S) (S = C or G) to the target ‘A’ nucleotide. Both SpCas9-evoA-BE4max and xCas9-evoA-BE4max preferred a T<u>C</u> motif, but the other three CBEs more favored a C<u>C</u> motif (Fig. 2b). In addition, the third nucleotide either upstream (G or C at position −3) or downstream (G or C at position +3) to the target ‘C’ also contributed significantly to the editing efficiencies of CBEs (Fig. 2b).

**Fig. 2.**
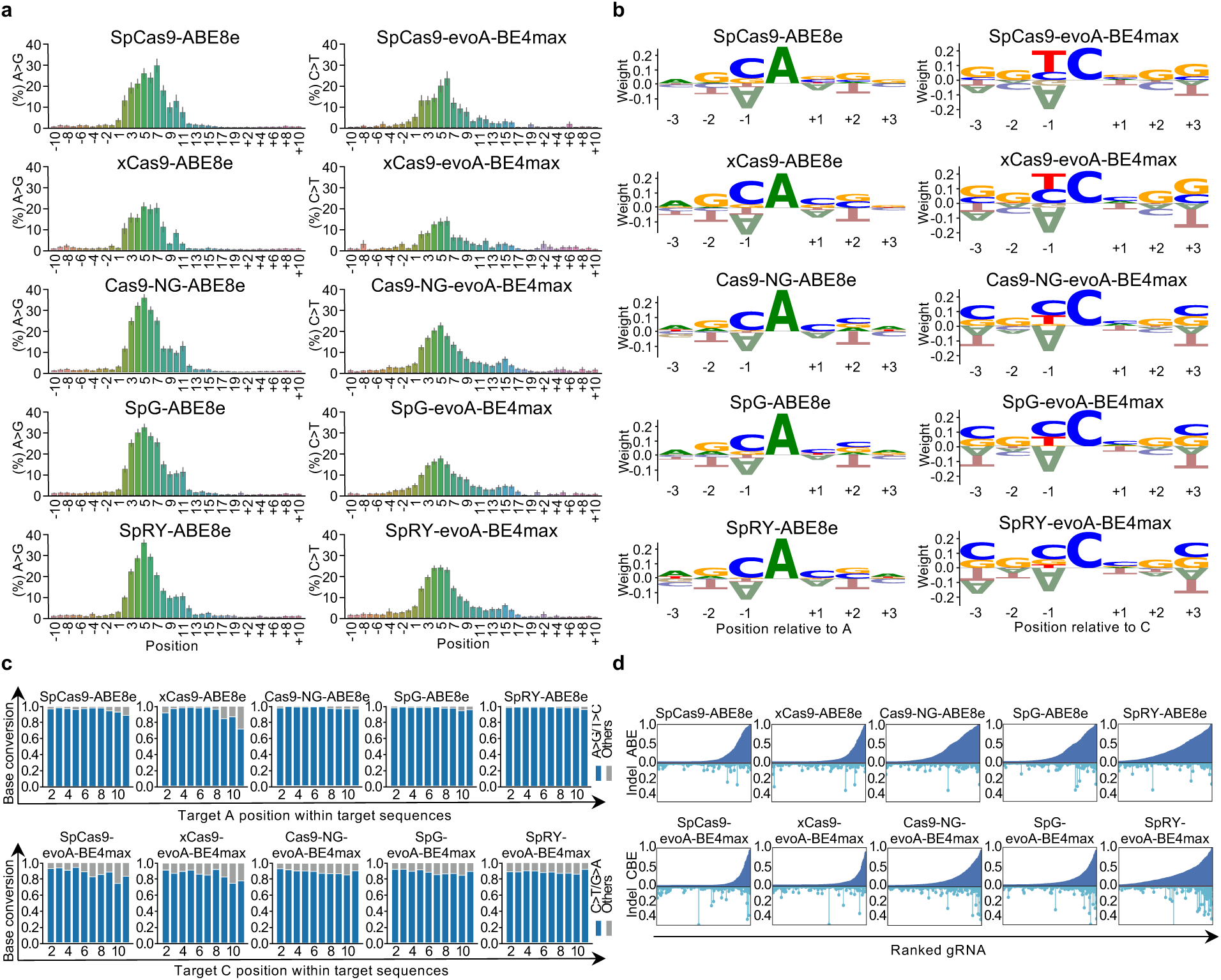
Characterization of editing efficiency, context, purity, and indel. **a**, Bar plot showing the editing efficiency (counting both default strand and complementary strand editing together) at different position. ‘-’ means 5’ end extension and ‘+’ is 3’ end extension to spacer. The error bar is standard error (SE). **b**, The nearby nucleotide preference of strong editing activity events (editing efficiency > 0.05). The y-axis is normalized position-specific score. The x-axis is relative position to intended editing nucleotide. The central ‘A’ or ‘C’ is the target position. **c**, The edited base purity for indicated BE at different editing position. Blue bar is default A>G/C>T base editing, and the gray bar shows the percentage of all other editing types. **d**, Relationship between base editing and indel. Top bar plot shows the ranked base editing, and the bottom lollipop is the associated indel frequency.

We next examined the editing purity (standard A•T-to-G•C or C•G-to-T•A conversion versus all possible nucleotide transitions at the target position for ABE or CBE, respectively) for each BE. In target windows between position 3 and 7, all ABEs showed high editing purity, whereas SpCas9-ABE8e and xCas9-ABE8e apparently exhibited less purity outside of the typical editing window, especially starting from position 9 and beyond (Fig. 2c). All the CBEs showed dramatic non-standard editing and less purity than ABEs, and such impurity generally displayed little positional preference, which was also different from that of ABEs (Fig. 2c). The sequence motifs for editing purity of ABEs generally resembled editing efficiency motifs that favored C<u>A</u> dinucleotide, but more weighting was observed on downstream sequence context with <u>A</u>WC (W = A or T) as a preference and <u>A</u>SW as an aversion (Extended Data Fig. 3a). For CBEs, unlike a favored Y<u>C</u> (Y = C or T) target observed in efficiency motif analysis, such dinucleotides were not observed in purity motifs. Rather, <u>C</u>C (for SpCas9-evoA-BE4max) or <u>C</u>S (for other CBEs) targets were more likely to have a pure C•G-to-T•A editing (Extended Data Fig. 3a). These data suggest that, when a high fidelity or purity of base editing is required, different preference of sequence context should be considered which cannot be directly obtained from simple efficiency model.

We further examined the unwanted indel outcomes accompanied by desired base editing. For all the BEs, the majority of high indel frequency events were associated with low base editing efficiency (Extended Data Fig. 3b), suggesting that the activities of indel generation and base conversion are generally uncoupled. However, some CBEs (especially SpRY-evoA-BE4max, Cas9-NG-evoA-BE4max and SpG-evoA-BE4max) tended to produce a broader and higher frequency of indels compared to other BEs (Extended Data Fig. 3b), and their top base editing events were associated with strongest indel activities (Fig. 2d), cautioning that a balance of desired base editing efficiency and indel frequency should be considered for these BEs.

### Targeting scope comparison for BEs

Although the nuclease activities and PAM preferences have been characterized for those Cas9 variants, how these BEs consisting of Cas9 nickase variants and base-converting enzymes determine their optimal targeting scope remains largely unknown and requires systematic characterization. Using the PAM-less gRNA tiling strategy and high throughput evaluation, we were able to analyze the base editing efficiencies of each BE for massive gRNAs with all possible PAM motifs. In Fig. 3a, we compared the editing activities of each BE corresponding to all possible 4-mer NNNN PAMs for base editing activities, which displayed dramatic differences between different BEs. For SpCas9-ABE8e and SpCas9-evoA-BE4max with classic Cas9 nickase, a general NGGN PAM was evident for these BEs, consistent with the NGG PAM for SpCas9 nuclease (Fig. 3a). However, a specific AGGT PAM was strongly disfavored for both SpCas9-ABE8e and SpCas9-evoA-BE4max, followed by CGGA aversion for SpCas9-evoA-BE4max. Except for NGGN, a second NAGN PAM block also showed some activities for SpCas9-ABE8e and, to a lesser extent, for SpCas9-evoA-BE4max. In addition, the fourth nucleotide ‘T’ within PAM was clearly disfavored for both BEs (Fig. 3a). For xCas9-conjugated ABE and CBE, a general NGNN PAM could be observed, but the activities of such PAM pattern were not very high, especially for xCas9-evoA-BE4max. The strongest PAM for xCas9-ABE8e was SGGY, while CGGT and TGGG were the top two PAMs for xCas9-evoA-BE4max. Cas9-NG- and SpG-derived BEs showed similar PAM patterns with apparently high activities for NGNN PAMs and relatively weaker NANN PAMs, especially for ABEs. Moreover, a strong GTGG PAM was observed for Cas9-NG- and SpG-derived ABEs but barely seen for CBEs, further demonstrating that the PAM preference of BEs is different from that of Cas9 nucleases and cannot be readily concluded from lessons of Cas9 variants. As expected, SpRY-derived BEs showed the broadest PAM patterns with appreciable base conversion activities. Both SpRY-ABE8e and SpRY-evoA-BE4max were nearly PAM-less with only a few specific PAMs (e.g., TAAB) (B = C or G or T) exhibiting no activities (Fig. 3a). We then plotted the base editing efficiencies for each BE by breaking down their targets in all possible 3-mer PAMs. Again, we saw the strongest PAM bias for SpCas9-derived BEs and the broadest PAM compatibility for SpRY-derived BEs (Fig. 3b). Taken together, these data comprehensively showed varied PAM characteristics and corresponding base editing efficiencies for each BE and are important to determine whether a target is editable by a given BE.

**Fig. 3.**
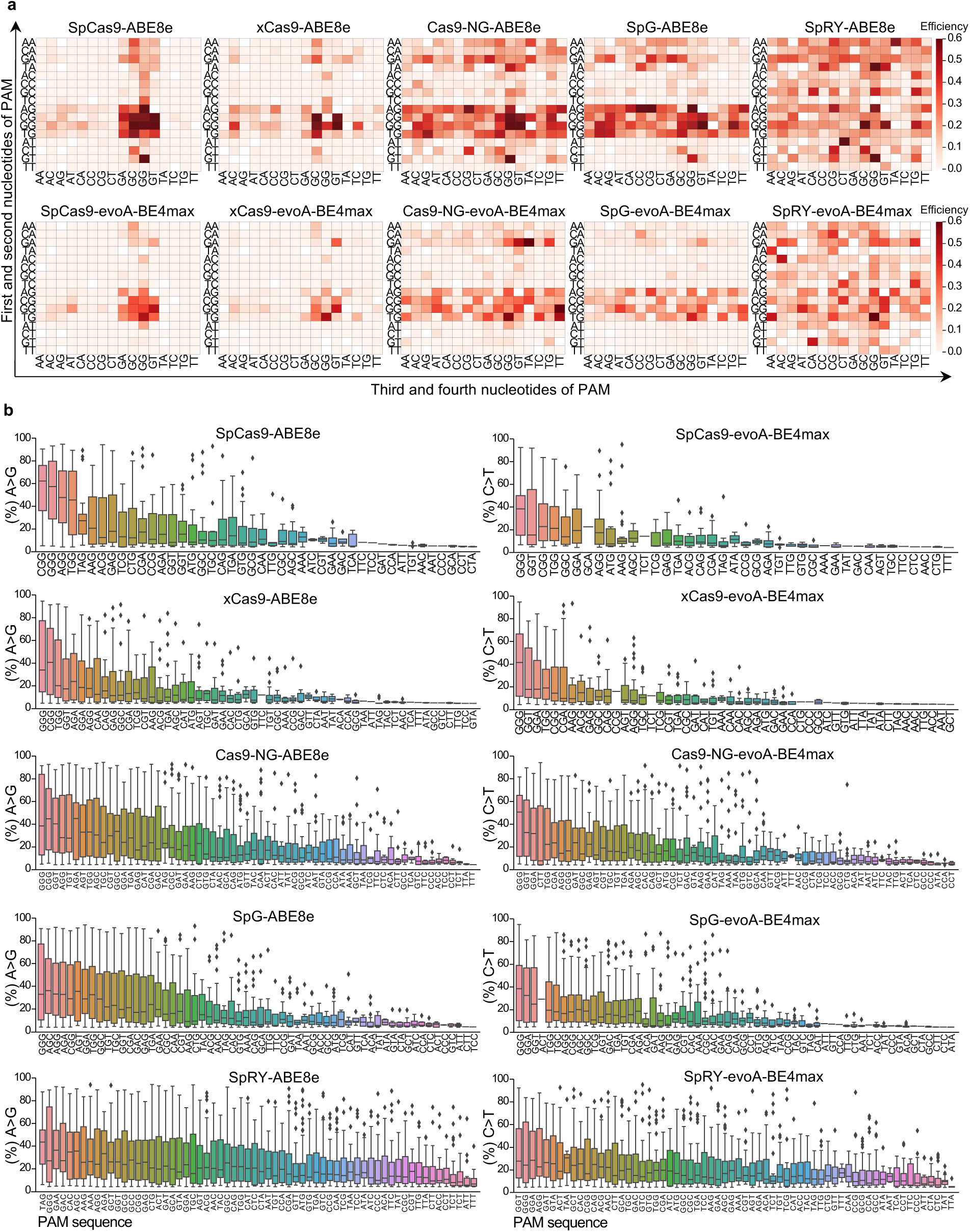
Characterization of PAM compatibilities for indicated BEs. **a**, The 4-nucleotide PAM patterns for each BE according to base editing activities. The y-axis is the first two-nucleotide, and the x-axis is the last two-nucleotide. **b**, The ranked PAM patterns in 3-nucleotide for each BE according to base editing activities. The box shows the interquartile range (IQR) from the 25th to the 75th percentile, with a line marking the median (50th percentile). Whiskers extend to 1.5 times the IQR.

### A deep learning model for predicting the editing efficiency and outcome of Bes

To facilitate the usage of these BEs and the choice of optimal BE, gRNA and target combinations, we developed a deep learning model, named BEEP (<u>B</u>ase <u>E</u>diting <u>E</u>fficiency <u>P</u>redictor), to inform the relationship between sequence and base editing efficiency by taking advantage of the above high-throughput evaluation data. We used a convolutional neural network (CNN) comprising three convolutional layers and three fully connected layers as BEEP’s base structure. The only input for the BEEP model is a 29-nt sequence, which includes a 19-bp protospacer and two 5-bp flanking sequences for adapting different spacer lengths and covering the PAM region (Fig. 4a). BEEP is designed to predict base editing efficiency at all possible positions within the spacer. For a specific editing position, the input sequence was converted into 3 sub-sequences with the same length: 1) target sequence, which is a copy of the input and contains the full information including the PAM sequence; 2) guide sequence, which highlights the spacer information and masks others; and 3) editing sequence which only contains the editing context (3 bp extended to both ends of the target editing) and is important to code the editing position (Fig. 4a). Each sub-sequence was encoded to a four-dimensional binary matrix by one-hot encoding. In addition, the Cas9 variant type was represented in a four-dimensional binary matrix with the same size. Finally, these 3 sub-sequence matrices and the Cas9 variant matrix were used to learn features by the CNN model and predict efficiency by the fully connected network (Fig. 4a). The output of BEEP is a score, named “BEEP Score” and ranging between 0 and 1, to indicate the base editing efficiency at the target position using a specific Cas9 variant. For the 29-nt input sequence, BEEP generates a list of all editing positions and Cas9 variant combinations with the predicted BEEP score.

**Fig. 4.**
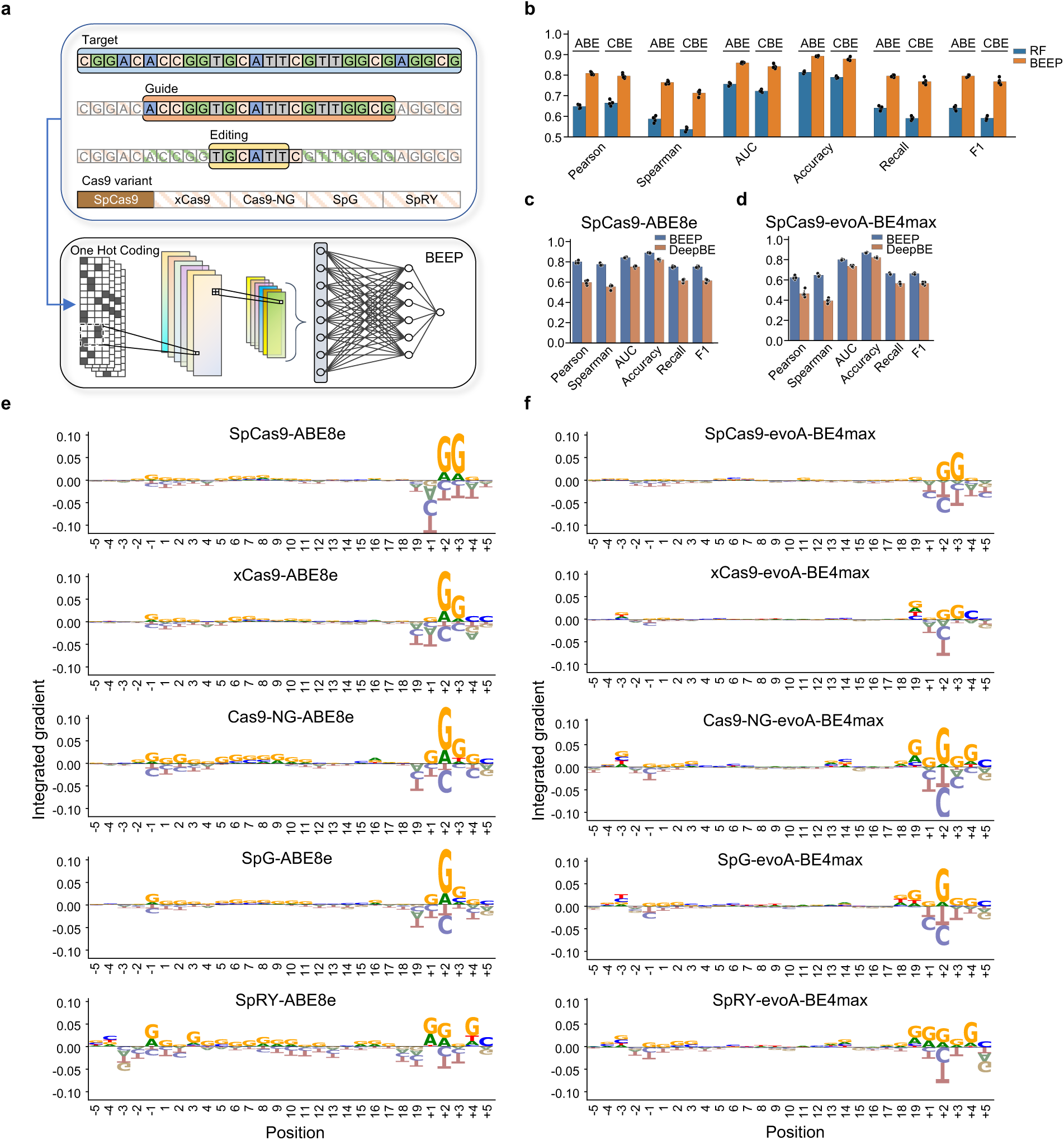
A machine learning model BEEP for predicting the base editing efficiencies and outcomes. **a**, The schematic diagram of BEEP model. Four feature vectors can be extracted from the extended target sequence and then one-hot encoding is applied to generate input matrix for the CNN model. **b**, Performance comparison of BEEP to traditional random forest (RF) model using 5 measure matric, including Pearman correlation, Spearman correlation, AUC (Area Under Curve), Accuracy, Recall and F1 score. The top quantile base editing events are treated as positive events. **c**,**d**, Performance comparison of BEEP to public deep learning model DeepBE for SpCas9-ABE8e (**c**) and SpCas9-evoA-BE4max (**d**), respectively. The scatters in the bar plot are generated from 5-fold cross-validation experiment. **e**,**f** Feature importance for base editing activity of ABEs (**e**) or CBEs (**f**) at each nucleotide position by Integrated Gradients (IG) analysis.

We compared BEEP with the Random Forest (RF) algorithm and a published deep learning-based prediction model, DeepBE^46^, using our tiling gRNA dataset (Supplementary Data 1). A matrix comprising the Pearson correlation, Spearman correlation, AUC, Accuracy, Recall, and F1 scores was built to evaluate the performance of different models using 5-fold cross-validation. For RF algorithms, the input feature matrix was generated manually based on the nucleotide type, editing position, GC content, melting temperature, and Cas9 variant type. For DeepBE, we predicted the efficiency using the default parameters and focused on SpCas9. BEEP not only showed better performance than RF (Fig. 4b), but also outperformed DeepBE (Figs. 4c,d) for both ABEs and CBEs in our BE toolbox. For practical usage, we further developed a web server of BEEP (http://beep.weililab.org/) to facilitate the end-users in selecting optimal BE-gRNA-target combinations (Extended Data Fig. 4).

To better understand the base editing preference, we applied the Integrated Gradients (IG), an explainable AI framework, to estimate the relationship between the BEEP model predictions and its associated features. Based on the output of IG, we calculated the preferences of sequence position and nucleotide composition in the input sequence. As shown in Figs. 4e,f, the PAM sequence was much more critical than the spacer sequence for all the BEs, and both the classic and non-canonical PAM patterns were identified via IG (Fig. 3a). Interestingly, in addition to 3’-end PAM, Cas9-NG-, SpG- and SpRY-derived BEs also exhibited weaker but noticeable sequence preferences at certain positions of 5’-distal region, especially upstream of the spacer cassette. In contrast, SpCas9- and xCas9-derived ABEs and CBEs barely showed 5’-end sequence preferences (Figs. 4e,f). These results further illuminate important sequence preferences other than PAM in determining the base editing outcomes for a given BE.

### Installment of disease-associated SNVs in ClinVar database

As a majority of disease-associated genetic variants are SNVs^51^, BEs are promising tools for the modeling and correction of these disease-associated SNVs. In addition to therapeutic correction, installing these SNVs in model systems is also fundamental in the functional and mechanistic studies of many human diseases. Moreover, the causal relationship between most of these SNVs and corresponding diseases has not been established or evaluated due to the lack of highly efficient techniques to install the SNVs in the genome. For 2,655,785 disease-associated SNVs in ClinVar database^51, 52^, 21.78% and 44.22% of them can be installed theoretically via the typical base conversions using ABE (for A>G and T>C) and CBE (for C>T and G>A), respectively (Fig. 5a). Further considering constraints from PAM compatibility and editing window (generally position 3-8), around 29%-35% of these ABE or CBE targets are editable via the spCas9-based BEs (with NGG PAM). When using other Cas9 variants with relaxed PAMs (NGN or NRN), the editable targets dramatically increased to 77% or 96% for ABE and 82% or 96% for CBE (Fig. 5b), respectively, highlighting the significance of enlarging the targeting scope via PAM extension. Using BEEP, we predicted the editability of theoretical ABE or CBE targets of ClinVar SNPs for each BE (Fig. 5c), where the editing percentage by individual BE was generally consistent with that of PAM analysis (Figs. 5b,c). Due to the contribution of non-classic PAMs, new properties endowed to BE rather than learned from Cas9 nucleases, the predicted editing percentage was all slightly higher for corresponding BE than that for targets with indicated PAMs (Fig. 5b,c). Notably, SpCas9-ABE8e was predicted to install many more targets than those predicted by just NGG PAM (38.44% vs. 28.94%), underscoring the importance of non-classic PAMs for these BEs.

**Fig. 5.**
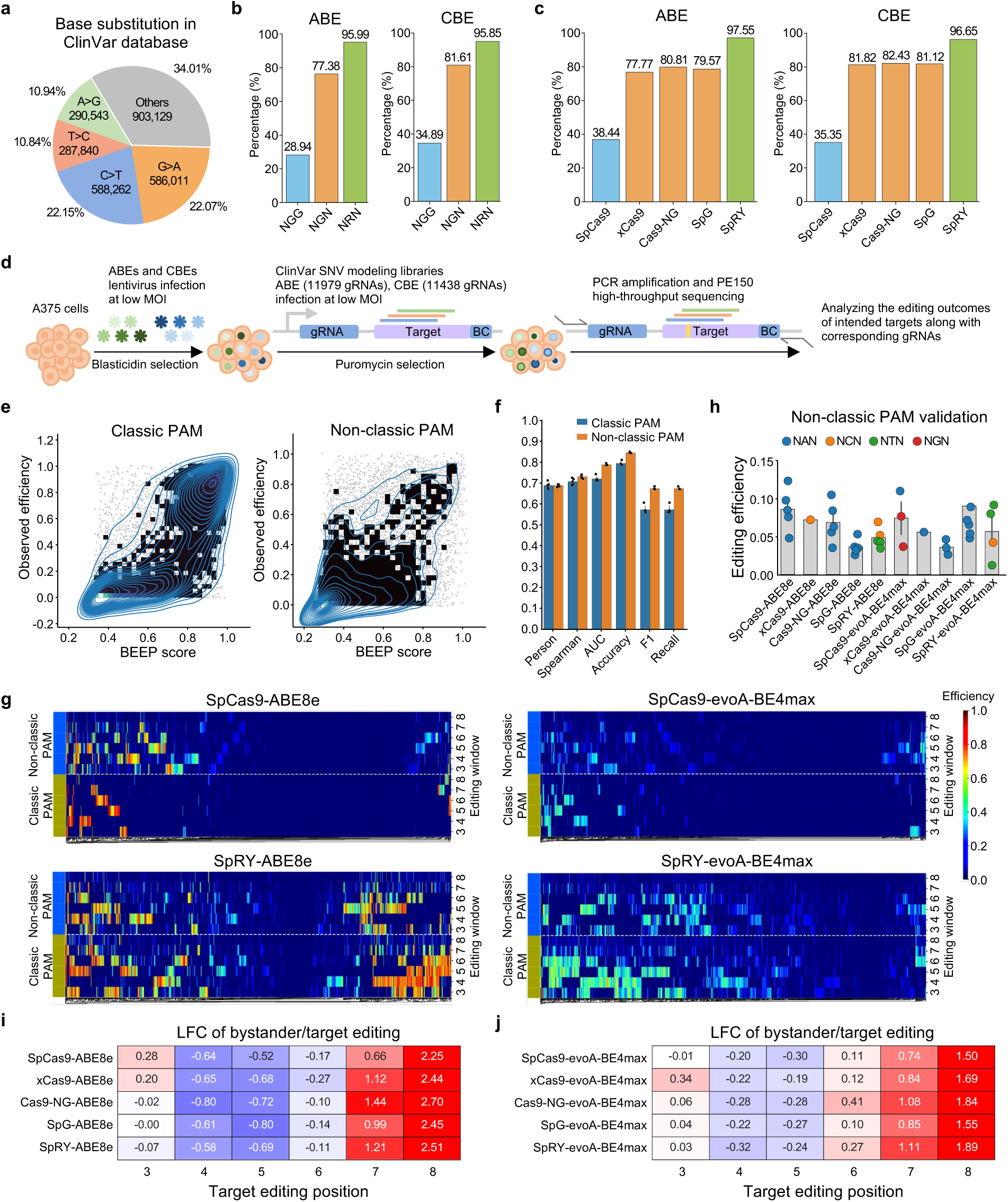
High-throughput installment of disease-associated SNVs in ClinVar database. **a**, Pie chart showing the statistics of base substitution types for ClinVar SNVs. **b**, The percentage of editable ClinVar SNVs by ABE/CBE with indicated PAM patterns within editing window position 3-8. **c**, BEEP-predicted editable ClinVar SNVs by indicated BEs when considering both classic and non-canonical PAM patterns. **d**, The schematic workflow to experimentally install ClinVar SNVs at scale using ABE and CBE libraries. **e**, The relationship between observed base editing efficiency and BEEP score when experimentally installing ClinVar SNVs with our BEs. **f**, The performance of BEEP prediction for base editing events with classic and non-classic PAMs using high-throughput ClinVar SNV modeling data. **g**, Heatmap showing the base editing performance of indicated BEs for targets with classic and non-classic PAMs in ClinVar SNV modeling data. **h**, Individual validation of base editing activities for targets with non-classic PAMs. Each dot indicates a target and PAM patterns are color-coded. **i**,**j**, The log2 fold change (LFC) value of bystander editing versus target editing across different positions for each ABE (**i**) or CBE (**j**).

To further evaluate the accuracy of our BEEP model and explore the efficacy of installing ClinVar SNVs using different BEs, we designed two additional libraries comprising 11,979 and 11,438 gRNA-target matched pairs to install 1,787 and 1,771 ClinVar SNVs by our ABEs and CBEs, respectively (Fig. 5d and Supplementary Data 2). Each SNV was covered by an average of 6 gRNAs without considering PAM restriction. The gRNA-target matched libraries were transduced lentivirally at a low MOI (∼0.3) into 10 lines of above BE expression-calibrated A375 cells followed by puromycin selection. After 7 days, the genomically integrated gRNA-target cassettes were isolated from cells and the editing outcomes of intended targets along with corresponding gRNAs were resolved by PE150 high-throughput sequencing (Fig. 5d and Supplementary Data 2). Using both screening datasets as independent evaluations, we further validated the performance of BEEP in predicting the base editing efficiencies and outcomes. For targets with classic PAMs, a strong correlation was found between BEEP predictions and experimentally confirmed efficiencies, especially for those with high predicted efficiencies (Fig. 5e). We also observed a slightly skewed distribution, where BEEP prediction scores were higher compared with observed editing efficiency in many loci (Fig. 5e), suggesting that other factors beyond sequence features might also play a role in determining the actual base editing efficiencies. For non-classic PAMs, the editing efficiencies were generally lower than targets with classic PAMs, but the correlation remains high between BEEP scores and the observed editing efficiencies (Fig. 5e). We also thoroughly evaluated BEEP performance using additional measurements, including Pearman correlation, Spearman correlation, AUC, Accuracy, Recall, and F1 score, further demonstrating the power of BEEP in predicting the efficiencies of installing ClinVar SNVs for sites with both classic and non-classic PAMs (Fig. 5f). Consistent with the BEEP prediction (Fig. 5c), these BEs altogether can efficiently install 76.1% and 72.3% of intended SNVs in our designed ABE and CBE libraries with efficiency above 0.2, respectively (Extended Data Figs. 5a,b; Supplementary Data 2).

Importantly, many disease-associated SNVs without classic PAMs can be edited by these 10 BEs via non-classic PAMs (Fig. 5g and Extended Data Fig. 5c). To validate this, we lentivirally transduced individual gRNAs into BE-expressing A375 cells, amplified the targeted genomic region, and determined the base conversion status for multiple endogenous SNV sites. As shown in Fig. 5h, the editing efficiencies for most of the tested targets with non-classic PAMs are between 5%-10%. Notably, such editing efficiency tended to be underestimated as the expression of BE and gRNAs were kept minimal due to the use of low MOI. In real applications, the editing efficiency is expected to increase with enhanced expression of BE components. To provide end-users a direct and comprehensive reference in modeling ClinVar SNVs, we predicted 52,579,530 BE-gRNA-target combinations for installing 1,752,651 disease-associated SNVs with BEEP, whose results can be freely accessible and downloaded via http://beep.weililab.org/.

In addition to the intended target nucleotide editing (A or C for ABE or CBE, respectively), other undesired A or C nucleotides within the editing window could also be modified by the corresponding BEs. Such bystander editing matters, especially in the scenarios of precision base conversion. With these high-throughput ClinVar SNV installment data (Supplementary Data 2), we were able to evaluate the characteristics of bystander editing for these BEs. As shown in Figs. 5i,j, the bystander editing was significantly high when the target site was located at position 7 or 8 for both ABEs and CBEs. In contrast, for target site positioned at 4 or 5, the bystander editing was kept minimal. In particular, SpCas9-ABE8e, xCas9-ABE8e, and xCas9-evoA-BE4max showed higher bystander editing at target position 3 than other BEs, and ABEs generally displayed weaker bystander editing at target position 6 than CBEs. In addition to the target position, different PAM patterns also affected the variations of bystander editing (Extended Data Figs. 5d). These features of bystander editing are helpful, especially when precision base editing is required.

### Interrogating cancer-associated mutations for drivers underlying BRAF inhibitor resistance

In addition to modeling or correcting disease-associated SNVs, BEs are widely used for studying single nucleotide mutations in cancer. As a showcase application, we explored the application of BE in identifying functional mutations in melanoma that confer resistance to BRAF inhibitors. We designed an ABE screen library for installing 1,243 highly frequent A>G somatic mutations in melanoma with 4,511 gRNAs (Supplementary Data 3). We transduced this lentiviral library into either SpCas9-ABE8e- or SpRY-ABE8e-expressing A375 cells at a low MOI (∼0.3), followed by puromycin selection. The edited cell pool was further split into multiple groups treated with either vehicle control (DMSO) or BRAF inhibitors (Vemurafenib or PLX4720) for three weeks to allow the positive selection of drug-resistant cells (Fig. 6a; see Methods). The genomically integrated gRNAs were quantified by high-throughput sequencing and analyzed with the MAGeCK algorithm^53^ (Supplementary Data 3). The two biological replicates for each condition were highly correlated (average Pearson’s *r* = 0.94 across all the conditions) (Extended Data Fig. 6a), indicating a high reproducibility of these base editing screens. As expected, the control gRNAs intended to disrupt splicing sites of essential genes in A375 cells were negatively selected in vehicle conditions (Extended Data Fig. 6b). For drug resistance hits, there was a significant overlap between Vemurafenib and PLX4720 treatment groups for each ABE (Extended Data Fig. 6c), further highlighting the consistency of BE screens. On the other hand, the overlap between SpCas9-ABE8e and SpRY-ABE8e groups for either drug was not significant (Extended Data Fig. 6c), suggesting a differential target profile for the two ABEs. Moreover, among the top drug-resistant hits, both *NRAS*_c.182A>G and *KRAS*_c.182A>G consistently ranked top in SpRY-ABE8e screens, albeit with less pronounced selection in SpCas9-ABE8e screens (Fig. 6b). Considering a high prevalence of *NRAS* mutations instead of *KRAS* in melanoma^54, 55^, we decided to follow up with the *NRAS*_c.182A>G variant. The *NRAS*_c.182A>G mutation results in a Q61R amino acid substitution of human NRAS protein, which leads to a more oncogenic GTP-bound form in activating downstream pathways^56^. To validate the function of *NRAS*_c.182A>G mutation during BRAF inhibitor resistance, we individually introduced two independent gRNAs (gRNA-1 and −2) mediating such base conversion into SpRY-ABE8e-expressing A375 cells and monitored the base editing frequency of *NRAS*_c.182A>G event before and after BRAF inhibitor treatment. As expected, a significant enrichment of edited *NRAS*_c.182A>G allele was observed in target gRNA-expressing samples upon drug treatment versus DMSO control (Fig. 6c). Furthermore, the surviving cell number for those samples expressing target gRNAs in the presence of BRAF inhibitors was significantly higher than those control samples (Fig. 6d,e). In addition, we knocked out endogenous *NRAS* expression and then lentivirally transduced either PAM-mutated wild-type (WT) NRAS-WT or NRAS-Q61R variant to determine their responses to BRAF inhibitor treatment (Extended Data Figs. 6d,e). As shown in Fig. 6f, only NRAS-Q61R-expressing cells, but not NRAS-WT or vector control samples, could confer cellular resistance to BRAF inhibitors. These data demonstrate that *NRAS*_c.182A>G mutation alone is sufficient to induce melanoma cell resistance to BRAF inhibitors. Given that around 20-30% of melanomas are *NRAS*-mutant and the majority of NRAS mutations reside in the Q61 site^54, 55^, these findings suggest a potentially important biomarker to inform the clinical response to BRAF inhibitors.

**Fig. 6.**
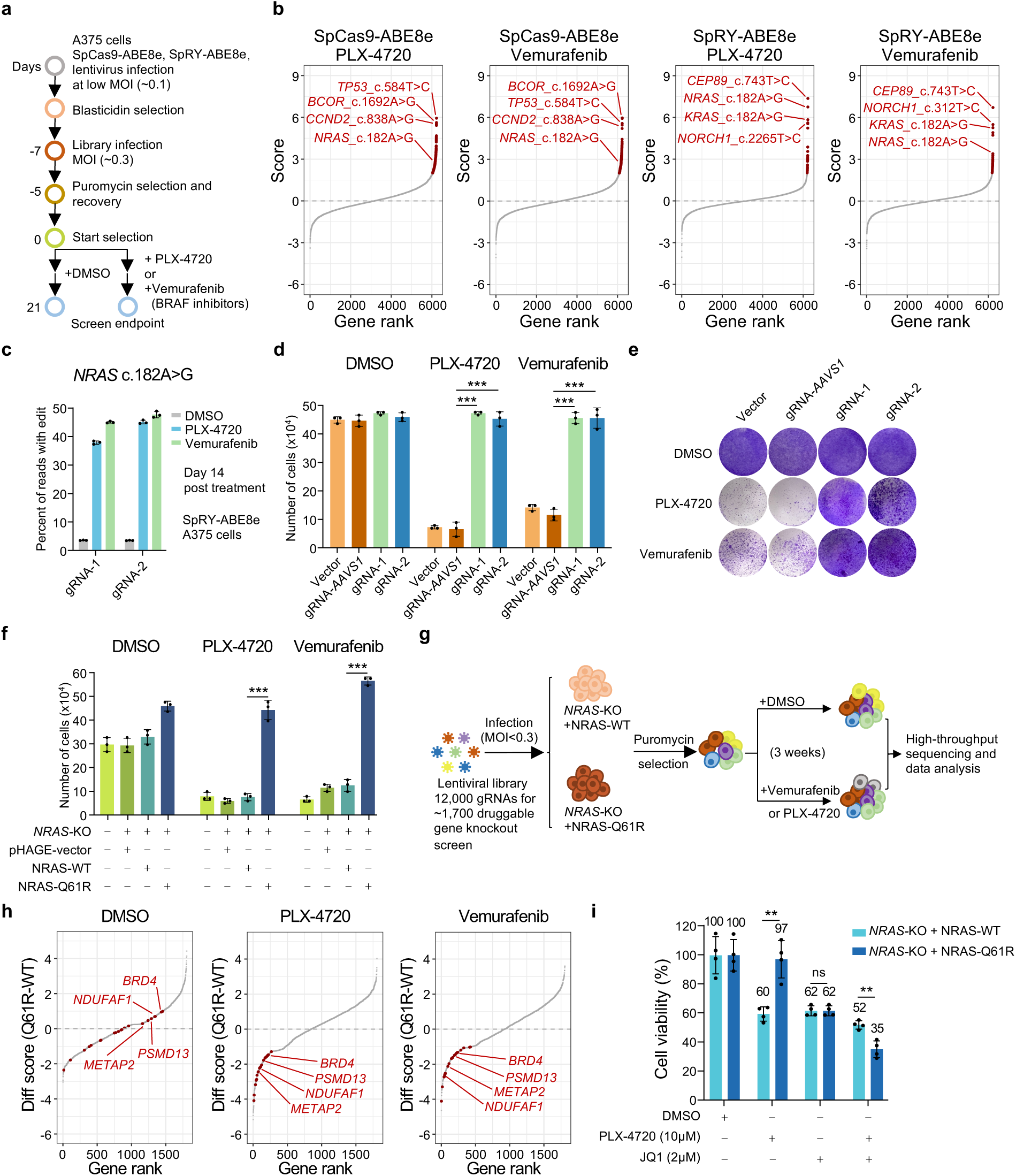
BE screening for driver mutations underlying BRAF inhibitor resistance in melanoma. **a**, Workflow of BE screening to identify driver mutations responsible for BRAF inhibitor (PLX-4720 or Vemurafenib) resistance in melanoma. **b**, The ranked list of interrogated gene mutations according to β-score in screens with indicated BEs and drug treatment. Representative hits are highlighted. **c**, Percentage of reads containing the intended edit (*NRAS* c.182A>G) for SpRY-ABE8e-expressing A375 cells lentivirally transduced with two independent gRNAs (gRNA-1 or −2) for base editing in the presence or absence of indicated drug treatment for 14 days. Mean ± SD with n = 3. **d**,**e**, Cell number quantification (**d**) or Crystal violet staining (**e**) of SpRY-ABE8e-expressing A375 cells lentivirally transduced with two independent gRNAs (gRNA-1 or −2) targeting *NRAS* c.182A>G after 14 days of DMSO or indicated drug treatment. Empty vector without specific gRNA and gRNA targeting *AAVS1* serves as control. Mean ± SD with n = 3 biological replicates. Unpaired two-sided *t* test, \*\*\**p* < 0.001. **f**, Replenishment of PAM-mutated NRAS-Q61R but not NRAS-WT into *NRAS* knockout single clonal A375 cells showed significant drug resistance to PLX-4720 or Vemurafenib treatment. Cell number was counted by a hemocytometer. Mean ± SD with n = 3 biological replicates. Unpaired two-sided *t* test, \*\*\**p* < 0.001. **g**, Workflow of druggable gene CRISPR knockout screens in NRAS-WT- and NRAS-Q61R-replenished *NRAS* knockout cells. **h**, The ranked list of druggable gene hits according to a differential β-score between NRAS-Q61R- and NRAS-WT-expressing cells under indicated conditions (DMSO, PLX-4720 or Vemurafenib treatment). Representative gene hits showing a preferential synthetic lethal effect with BRAF targeted drugs in resistant cells are highlighted. **i**, MTT assay showing the cell viability for NRAS-WT- or NRAS-Q61R-replenished *NRAS* knockout cells treated with PLX-4720 or JQ1 along or combination. Mean ± SD with n = 4 biological replicates. Unpaired two-sided *t* test, \*\**p* < 0.01. ns means not significant.

To further explore the potential therapeutic targets against NRAS-Q61R-induced resistance, we performed a CRISPR knockout screen using our druggable gene Cas9-gRNA library^57^ (around 12,000 gRNAs targeting ∼1,700 druggable genes) in both resistant (expressing NRAS-Q61R on top of *NRAS* knockout) and sensitive control (expressing NRAS-WT on top of *NRAS* knockout) cells in duplicates (Fig. 6g and Extended Data Fig. 6f; Supplementary Data 4). If a druggable gene was more negatively selected in resistant cells than in sensitive control cells, the inhibitory drugs targeting this gene might be potentially effective therapeutics against BRAF inhibitor-resistant melanoma. Indeed, we found several druggable genes that were apparently synthetic lethal with either BRAF inhibitor but not with DMSO control (Fig. 6h), including BRD4, a well-characterized transcriptional and epigenetic regulator as well as a promising drug target for blood and solid malignancies^58–60^. Indeed, the BRD4 inhibitor JQ1^61^ and BRAF inhibitor PLX-4720 displayed a significantly synergistic effect in reducing the growth of resistant NRAS-Q61R-expressing cells (Fig. 6i), suggesting JQ1 as a candidate drug to overcome BRAF inhibitor resistance in melanoma. Altogether, these data showcase the power and tentative strategies of applying our BE tools and other techniques to functionally characterize cancer mutations at scale.

## Discussion

Here we systematically benchmarked the performance of several ABEs and CBEs which altogether can theoretically cover almost the most targets for A•T-to-G•C or C•G-to-T•A conversion with high efficiency. These BE tools exhibited variable but collectively good performance in installing SNVs of ClinVar database or point mutations in cancers. The deep learning model BEEP, trained on the high-throughput benchmark data, accurately predicts the editing efficiency and outcomes for these BEs. We further built a freely accessible webserver for end-users to prioritize appropriate BE/gRNA combinations for the target of interest.

Many progresses have been made to develop new or improved BE tools with specific features to meet different purposes^62, 63^. These new technologies greatly expanded the available tools that can be used, but also pose challenges for users to determine suitable BEs for different applications. Among many features of BEs, the editing efficiency and the targeting scope are the two major factors considered in the first place, as they largely determine whether a target site can be edited with high efficiency. To meet the editing needs for as many target sites as possible, we combined highly efficient deaminase domains with multiple Cas9 nickase variants bearing different PAM preferences. Notably, other BE properties such as context combability, editing window, fidelity, and off-target effect also contribute to the final editing outcomes and affect the choice of suitable BEs. In general, the set of ABE/CBEs we benchmarked here could serve as the first lines of choice for most of the A•T-to-G•C or C•G-to-T•A conversion purposes. Calibrated BE expression, long targets with uniform context, and the same tiling gRNA libraries in our study allowed us to systematically benchmark different BEs in a comprehensive and unbiased manner. We revealed detailed and different characteristics of these BEs regarding editing efficiency, outcome, purity, context, indel, PAM, bystander editing, etc. Such comparative analysis better informs the users to select more suitable BEs for the targets of interest. The derived BEEP model performed better than existing models in predicting the editing efficiency and outcomes. The freely accessible BEEP web server further enables users to easily select optimal BE-gRNA combinations for a given input target. Using these BEs, we experimentally installed 3,558 disease-associated SNVs in ClinVar dataset, and 20.1% of them would not be edited if just considering the classical PAM restriction. More importantly, using BEEP model, we systematically predicted candidate BE-gRNA combinations for all 1,752,651 ClinVar SNVs, providing a critical resource for the community. We also utilized these BEs to interrogate the function of cancer-associated point mutations. The combination of base editing screens and the traditional gene knockout screens not only reveals the driver mutations underlying BRAF inhibitor resistance, but also provides a roadmap to identifying actionable targets to overcome drug resistance.

In summary, our study provides a set of thoroughly evaluated ABEs/CBEs to edit A•T-to-G•C or C•G-to-T•A with high editing efficiency and broad targeting scope. The deep learning model BEEP further guides users to better select suitable BE-gRNA pairs for a target of interest. We expect that our efforts can serve as a starting point or guidance for general users to unleash the power of base editing in broader scenarios. Such evaluation or investigation strategy could also be extended to fully characterize or benchmark other types of BEs.

## Methods

### Plasmid construction

To construct the base editor plasmids, the pLenti-Cas9-Blast (Addgene, 52962) vector was digested with XbaI (Takara, 1634) and BamHI (Takara, 1605) restriction enzymes to remove the Cas9-encoding sequence and replaced with synthesized (Synbio Tech) human codon-optimized (GenScript scoring algorithm) TadA-8e, evoAPOBEC1, UGI fragments and overlapping-PCR mutated Cas9 variants through Gibson Assembly (TIANGEN, 4992813). The following BE plasmids were constructed: pLenti-SpCas9-ABE8e-Blast, pLenti-xCas9-ABE8e-Blast, pLenti-Cas9-NG-ABE8e-Blast, pLenti-SpG-ABE8e-Blast, pLenti-SpRY-ABE8e-Blast; pLenti-SpCas9-evoA-BE4max-Blast, pLenti-xCas9-evoA-BE4max-Blast, pLenti-Cas9-NG-evoA-BE4max-Blast, pLenti-SpG-evoA-BE4max-Blast, and pLenti-SpRY-evoA-BE4max-Blast. The detailed sequence information can be found in Supplementary Note 1.

To generate the gRNA-expressing plasmid (pLenti-CRISPRv2-Guide-Puro) for high-throughput screening assays and single site validation, the pLenti-CRISPRv2 plasmid (Addgene, 52961) was digested with XbaI and BamHI restriction enzymes to remove the Cas9-encoding sequence while keeping the puromycin resistance gene and the gRNA-expressing cassette. The libraries and individual gRNA sequences inserted into this vector were listed in Supplementary Data 1,2,3,5.

For the preparation of pHAGE-NRAS-WT-Blast or pHAGE-NRAS-Q61R-Blast expression plasmid, the pHAGE-EF1α-Blast vector was digested with XbaI and BamHI and then ligated with the PCR amplified NRAS coding sequence (NM_002524) or variant sequence (*NRAS* c.182A>G, p.Q61R) by T4 DNA ligase.

### Oligonucleotide library design

For high throughput BE evaluation assays, we designed tiling gRNAs regardless of PAM restriction on EGFP and firefly luciferase coding sequences. The gRNA-barcode oligo fragments were firstly array-synthesized and then cloned into the plasmid vector (pLenti-CRISPRv2-Guide-Puro). The longer target fragments of EGFP and luciferase were then inserted between the gRNA cassette and random 15 bp barcode (Extended Data Fig. 1d). To simultaneously sequencing the whole fragments encompassing the gRNA cassette and the corresponding target region, the EGFP and luciferase target sequences were split into six fragments (with each fragment ∼350 bp long) to accommodate the length limit of PE250 sequencing mode. Therefore, the evaluation library was divided into six sub-libraries named EGFP1, EGFP2, luciferase1, luciferase2, luciferase3 and luciferase4. A total of 3404 gRNA-target-barcode combinations were included in the final evaluation library (Supplementary Data 1).

For ClinVar SNV modeling library design, disease-related mutations were downloaded from NCBI ClinVar website with classification in Pathogenic, Likely Pathogenic or Uncertain Significance. We made sure the target editing position within the editing windows and scanned all possible guides for individual mutation without considering PAM restriction. A total of 11,979 and 11,438 gRNA-target matched pairs were designed to cover 1,787 and 1,771 target sites of ClinVar SNVs for ABE and CBE, respectively (Supplementary Data 2). To achieve a high editing performance, the target SNV is always in the editing window (position 3-8) for corresponding gRNAs. The target fragment was 50 bp long by extending 15nt on both 5’ and 3’ ends of the guide (gRNA-matching sequence). An average of 6 gRNAs were placed for each target nucleotide.

For drug resistance screens, we designed a library for ABEs to model those point mutations of high frequency in human melanoma and explored their effects on BRAF inhibitor sensitivity. This ABE library contained 6,229 gRNAs with 4,511 gRNAs targeting 1,243 highly frequent point mutations (>1.5% mutated genes and top 100 frequently mutated variants of all melanomas in COSMIC database^64^) to install A>G mutations by ABEs. The rest of gRNAs were designed to target the splicing sites of general essential genes and the *AAVS1* safe harbor locus. The detailed information could be found in Supplementary Data 3.

### Plasmid library construction

To construct plasmid library for high throughput BE evaluation, a two-step protocol was deployed. *Step I: Generation of the first plasmid library containing pairs of gRNA sequences and corresponding barcodes.* The pLenti-CRISPRv2-Guide-Puro vector was digested with BsmBI (New England Biolabs, R0739L) and NheI (Takara, 1622), and the six target oligo libraries were amplified by PCR (introducing overlapping sequence with vector). The vector and PCR-amplified oligo libraries were gel purified using a GeneJET Gel Extraction Kit (Thermo Fisher Scientific, K0692), and assembled using an 2X Gibson Assembly Mix at 50 °C for 1 h. After incubation, the mix was transformed into self-prepared electrocompetent Stable *E. coli* cells (New England Biolabs, C3040) by electro-transformation to reach the efficiency with at least 50X coverage representation of each clone in the designed library. The transformed bacteria were then cultured in liquid LB medium with Ampicillin for 16∼20 h at 30°C. The library plasmids were then extracted with Endo-Free Maxi-prep Plasmid Kit (TIANGEN, 4992194). *Step II: Generation of the second plasmid library by insertion of gRNA scaffold and target fragments.* The first plasmid libraries described above were digested with Esp3I (BsmBI) restriction enzyme (Thermo Fisher Scientific, FD0454) at 37 °C for 1.5 h. The linearized and dephosphorylated plasmids were separated on a 1.0 % agarose gel and purified using a GeneJET Gel Extraction Kit. The six target sequences with gRNA scaffold ahead were directly synthesized (Synbio Tech) and PCR amplified with Q5 High-Fidelity DNA polymerase (New England Biolabs, M0491L) using primers containing BsmBI restriction sites. These fragments were then cloned into the first plasmid libraries by T4 DNA ligase (Thermo Fisher Scientific, EL0016). The products were transformed into self-prepared electrocompetent Stable *E. coli* cells by electro-transformation to reach the efficiency with at least 50X coverage. The transformed bacteria were then cultured in liquid LB medium with Ampicillin for 16∼20 h at 30°C. The plasmid libraries were extracted with Endo-Free Maxi-prep Plasmid Kit and pooled together as the final BE evaluation plasmid library.

For ClinVar SNV modeling plasmid library construction, a similar two-step protocol was deployed. The first plasmid library was generated containing designed combinations of gRNA sequences, corresponding target sequences and barcodes. The gRNA scaffold sequence was then inserted via BsmBI site by T4 DNA ligase to get the final plasmid library. The construction of drug resistance BE screen plasmid library also followed similar procedures except that the first plasmid library only contains designed gRNA sequences without corresponding target fragment.

### Cell culture

HEK293FT, A375 (human melanoma cell line) cells were obtained from American Type Culture Collection (ATCC). All cells were maintained in DMEM (VivaCell BIOSCIENCES, C3114-0500) supplemented with 10% fetal bovine serum (FBS) (ExCell, FSP500) and 1% penicillin–streptomycin (VivaCell BIOSCIENCES, C3420-0100) and cultured with 5% CO_2_ at 37 °C. All cells were checked to ensure that they were free of mycoplasma contamination.

### Lentivirus production

HEK293FT cells were seeded in 100-mm culture dishes and transfected at 60-80% confluency. Opti-MEM reduced-serum medium (Gibco, 51200038) was mixed with 50 μL of Lipofectamine^TM^ 2000 Transfection Reagent (Invitrogen, 11668019), 6 μg of lentiviral vector, 4.5 μg of pCMVR8.74 and 3 μg of pMD2.G for a final volume of 0.5ml, incubated at room temperature for 20 min, and then added to the cell culture medium. After 12 h of transfection, the medium was replaced with fresh DMEM supplemented with 10% FBS. The lentivirus-containing medium was harvested after 48 h of transfection and centrifuged at 3,000 rpm for 5 min to remove the cell debris. The supernatant was divided into aliquots and kept frozen at −80 °C until use.

### High throughput BE evaluation assay

We firstly infected 5×10^5^ A375 cells with lentiviruses expressing ten different BEs (SpCas9-ABE8e, xCas9-ABE8e, Cas9-NG-ABE8e, SpG-ABE8e, and SpRY-ABE8e; SpCas9-evoA-BE4max, xCas9-evoA-BE4max, Cas9-NG-evoA-BE4max, SpG-evoA-BE4max, and SpRY-evoA-BE4max) at a low MOI (∼0.1) followed by 6 μg/mL blasticidin (Solarbio, B9300) selection for 7 days. These cells were further expanded before use. By using a low MOI, the expression of different BEs was kind of calibrated and kept equally minimal across samples. These BE-expressing cells were used not only for high-throughput assays but also for individual site evaluation in this study. For each BE evaluation, 6×10^6^ BE-expressing cells were then infected with lentivirus pool of BE evaluation library at a low MOI (∼0.3) in the presence of 10 μg/mL of polybrene (Solarbio, H8761) for 48 h before proceeding to 2 μg/mL puromycin selection for three days. Cells was then placed into refresh medium and cultured for additional two days before collecting the samples. Two biological replicates were performed for each condition. The genomic DNA was extracted from these cellular samples using the following steps. Add 1 mL Lysis buffer (300 mM NaCl, 0.2% SDS, 2 mM EDTA, 10 mM Tris-HCl pH 8.0) and 1 μL RNase A (100 mg/mL) in 15 mL conical tube containing the cellular pellet followed by mixing and incubation at 65°C for 1 h. Then add 10 µL Proteinase K (10 mg/mL) to continue incubation at 55°C for 6 h or overnight. Mix with 1 mL phenol/chloroform/isoamyl alcohol solution (25:24:1) and centrifuge at 12000 rpm for 10 min before taking the upper phase. The genomic DNA was precipitated by isopropanol and washed by 75% ethanol before dissolving in nuclease-free water. These genomic DNA and plasmid library DNA were used for constructing sequencing libraries. For the first round of PCR, a 100 µL reaction was performed, with 6–8 µg genomic DNA (maximum 10 µg per reaction) in each reaction, using Q5 High-Fidelity DNA polymerase with the primers listed in Supplementary Data 5. The resulting amplicons were combined before proceeding to the next step. A second round of PCR aims to attach Illumina adaptors and barcode samples. The second round of PCR was performed in a 100 µL reaction volume using 1 µL of the product from the first round of PCR for 10–12 cycles with the primers listed in Supplementary Data 5. These PCR products were gel purified and pooled for high-throughput deep sequencing using an Illumina PE250 sequencing mode (Novogene, China) to determine the editing outcomes.

### High-throughput installment of ClinVar SNVs

A375 cells (∼6×10^6^ for each group) were infected with lentiviruses expressing ten different BEs (SpCas9-ABE8e, xCas9-ABE8e, Cas9-NG-ABE8e, SpG-ABE8e, and SpRY-ABE8e; SpCas9-evoA-BE4max, xCas9-evoA-BE4max, Cas9-NG-evoA-BE4max, SpG-evoA-BE4max, and SpRY-evoA-BE4max) at a low MOI (∼0.1) followed by 6 μg/mL blasticidin (Solarbio, B9300) selection for 7 days. For each group, a total of 6×10^6^ BE-expressing cells were infected with lentivirus pool of ClinVar SNV installment libraries (either for ABEs or for CBEs) at a low MOI (∼0.3) in the presence of 10 μg/mL of polybrene (Solarbio, H8761) for 48 h before proceeding to 2 μg/mL puromycin (Solarbio, P8230) selection for 3 days. Cells was then placed into refresh medium and cultured for additional 2 days before collecting the samples. Two biological replicates were performed for each condition. Around 3.6×10^6^ (∼300x coverage) cells were collected for genomic DNA extraction. The gRNA-target paired fragment was then PCR-amplified to construct sequencing libraries. High-throughput sequencing (Illumina PE150 mode) was performed (Novogene, China) to determine the editing outcomes.

### Drug resistance BE screens

A total of 6×10^6^ SpCas9-ABE8e- or SpRY-ABE8e-expressing A375 cells were infected with lentiviruses of the drug resistance BE library at a low MOI (∼0.3) and seeded into two of 15 cm dishes. Two days later, the infected cells were selected with medium containing 2 μg/mL puromycin for 3 days followed by a two-day recovery in fresh medium. At least 2×10^6^ (∼300x coverage) cells were collected as the sample of Day 0, and the remaining cells (2×10^6^ cells for each group) were divided into three treatment groups: DMSO vehicle, 2μM PLX-4720 (Meilunbio, MB5133) and 2μM Vemurafenib (Meilunbio, MB2102). After 21 days, at least 2×10^6^ cells were collected as the sample of end time. The genomic DNA of all the collected samples was isolated and the gRNA fragment was PCR-amplified. High-throughput sequencing (PE150) was performed (GenePlus, China) to determine the abundance of the gRNAs. Two biological replicates were performed for each screen.

### Screening for genetic vulnerabilities of BRAF inhibitor-resistant cells

Around 6×10^6^ NRAS-WT- or NRAS-Q61R-expressing single clonal A375 cells were infected with a druggable gene Cas9-gRNA library^57^ at a low MOI (∼0.3) and seeded into 2×15 cm dishes for each group. Two days later, the infected cells were selected with 2 μg/mL puromycin for 3 days followed by a two-day recovery with normal medium. At least 3.6×10^6^ (∼300x coverage) cells were collected as Day 0 sample, and the remaining cells were divided into either DMSO vehicle or drug treatment (2μM PLX-4720 or 2μM Vemurafenib) groups. Each group contains at least 3.6×10^6^ cells at Day 0. After three weeks, the cells were collected as the samples of screening endpoint. The genomic DNA was isolated for all the collected samples and the gRNA fragment was PCR-amplified. The abundance of the gRNAs was determined by high-throughput sequencing with PE150 mode. JQ1 (Meilunbio, MB7243) was used as a BRD4 inhibitor.

### Western blot analysis

Collected cells were lysed in RIPA lysis buffer (Beyotime, P0013B) on ice for 20 min. The protein supernatant was collected by centrifuging at 14,000 rpm for 10 min at 4°C and mixed with 5x loading buffer. After boiling at 100 °C for 5 min, the samples were analyzed by SDS-PAGE and blotted with indicated primary and secondary antibodies. The primary antibodies were anti-FLAG (Invitrogen, MA1-91878), anti-NRAS (Santa Cruz Biotechnology, sc-31) and anti-GAPDH (Santa Cruz Biotechnology, sc-32233). The goat anti-mouse IgG (Santa Cruz Biotechnology, sc-516102) was used as secondary antibody.

### Editing efficiency measurement at endogenous sites

Individual gRNAs were designed and cloned into pLenti-CRISPRv2-Guide-Puro vector to measure the editing frequency of corresponding BEs at endogenous sites. Lentiviruses expressing these gRNAs were produced and infected the above BE expression-calibrated A375 cells at a low MOI (∼0.1) for 48 h. Cells were then selected with 2 μg/mL of puromycin for three days. After additional culture for two days, cells were collected to extract genomic DNA. The target regions were PCR amplified and subjected to deep sequencing on Illumina PE150 sequencing platform for determining the editing outcomes. The oligos, primers and gRNAs used in this section are listed in Supplementary Data 5.

### Computational analysis for high throughput BE evaluation

Customized Python script was used to scan the pair-end next generation sequencing (NGS) data read by read in the following steps: 1) identify spacer and barcode sequences from the R1 and R2 reads in the pair-end NGS data, respectively; 2) filter out any spacer-barcode mismatched reads and mark them as uncoupling reads; 3) locate target sequences from NGS reads based on the nearby fixed oligo components; and 4) extract and save target sequences into small fastq files based on the unique spacer or barcode. Next, CRISPResso2^65^ was used to analyze the base editing outcomes of these saved small fastq files with parameters: min_bp_quality_or_N = 30 and quantification_window_size = 10. In order to get confident analysis results, any small fastq files that the final used reads in CRISPResso2 analysis less than 200 will be filtered out due to the low qualified read count. We tried to minimize the recombination or uncoupling rates for the long reads by optimizing the experimental procedures (e.g., reducing the PCR cycles and using better DNA polymerase with lower recombination rates after extensive testing of multiple ones). The uncoupling rates for all the samples were between 55.2% and 70.1% with the median of 60.8%. Considering the lengthy fragment in this assay, such uncoupling rate is acceptable. We made sure enough read coverage in the analysis after removing these uncoupling reads.

### Editing and purity motif statistics

High editing (efficiency > 0.1) and high purity (purity > 0.9) activities were used to detect motifs nearby the editing events. We extended 3 bp on both ends of the base editing locus as the nearby sequence. Bio.motifs package were used to calculate nucleotide count at each position. We assumed that the nucleotide frequency is even and the default fraction should be 0.25. Then the weight was calculated as the difference to 0.25 and a positive weight means base preference. The weight was visualized by logomarker package as a motif.

### Computational analysis for high throughput ClinVar SNV modeling data

Similar to previous tiling screens, a customized python script was used to preprocess the NGS data and CRISPResso2 was used to analyze the base editing outcomes. As a comparison to BEEP, a Random Forest (RF) based machine learning method was used to predict the editing efficiency with default parameters. The ClinVar SNV modeling library was composed of 19 bp of gRNA sequences, whereas the DeepBE model was trained with 20 bp of gRNA or guide sequences. To address this, we extended the 19 bp guides by adding one base at either the 5’ or 3’ end of the sequence to form 20 bp guides. For each guide, we generated predictions by extending it in both directions (5’ and 3’), then calculated the average of the resulting prediction scores. This ensures that the predictions remain consistent with the DeepBE model’s training setup, while accounting for the slight discrepancy in guide length. A matrix composed by Pearman correlation, Spearman correlation, AUC, Accuracy, Recall and F1 score was used to evaluate the performance between BEEP and RF/DeepBE.

### Computational analysis for drug resistance BE screening data

The gRNAs were quantified by MAGeCK^53^ (version 0.5.9.2) “count” module with total read count normalization from raw fastq files. For gRNA level enrichment analysis, the gRNA counts of Day 21 samples were compared to those of corresponding Day 0 samples by “mle” module to obtain a β-score for each gRNA. A positive β-score indicates positive selection whereas a negative β-score means negative selection. The gRNAs targeting splice sites of top essential genes in A375 cells (with gene effect < −3 in DepMap; https://depmap.org/portal/) were employed as negative selection controls. For drug resistance positive selection, the hits were called as drug-resistant variants when the target gRNA bears a mean β-score > 2 across the two biological replicates.

### Computational analysis for druggable gene hits of BRAF inhibitor-resistant cells

MAGeCK^53^ (version 0.5.9.2) was employed for gRNA quantification and gene hit identification before and after the screening process. Raw fastq files were aligned to the druggable gene Cas9-gRNA library^57^ using the “count” module to obtain the gRNA counts. Normalized gRNA counts were used to calculate the β-score for gene level analysis with “mle” module. Each sample at the screening endpoint was compared with corresponding Day 0 sample. A positive β-score of a gene indicates positive selection whereas a negative β-score means negative selection. The original β-score for each sample was *Z*-score normalized and the mean value between two biologically replicates was assigned to each gene. To identify the synthetic lethal druggable gene hits with BRAF inhibitors preferentially in resistant cells, a differential (Δ) β-score was firstly calculated between the samples of NRAS-Q61R and NRAS-WT on the same condition (DMSO, PLX-4720 or Vemurafenib). The candidate hits were then defined with the following cutoff: (Δβ^PLX-4720^ and Δβ^Vemurafenib^) < −0.5 & (Δβ^PLX-4720/Vemurafenib^ – Δβ^Vehicle)^ < - 0.5 & (β^Q61R-PLX-4720^ and β^Q61R-Vemurafenib^) < −1. The detailed information can be found in Supplementary Data 4.

### BEEP deep learning model development

The input required by the BEEP model is an extended spacer sequence (5 nt extended in both 5’ and 3’ ends). The extension is critical to the prediction performance as it can cover the PAM region which plays an important role in the CRISPR/Cas system. BEEP will generate 4 vectors based on the extended sequences: 1) target vector - it’s the original extended sequence; 2) guide vector – it’s the spacer sequence and any other nucleotides will be masked; 3) editing vector – it’s used to highlight the editing position and extend 3 nt in both ends; and 4) Cas9 variant – it’s used to represent different Cas9 variants. All the 4 vectors are of the same length and this fixed length is a key to show relative location information. Next, these vectors will be converted into integer format using one-hot encoding and the masked nucleotides will be encoded as zero-vector. A Convolutional Neural Network (CNN) is used to extract the spatial features from the encoded input matrices and then a Fully Connected Neural Network is used for the final prediction. A BEEP Score for each guide will be output to indicate the predicted editing efficiency. The higher the BEEP Score is, the more likely the guide is to be effective. During the hyperparameter tuning process, we focused on four parameters: the number of units in the dense layer, the dropout rate, the optimizers, and the learning rate. For simplicity, each parameter was treated as a discrete value and a grid search was used to try all combinations of these discrete parameters. The Mean Squared Error (MSE) of the model at every epoch was tracked to determine the best combination of hyperparameters.

### Deep learning model interpretability

Integrated Gradients (IG) is implemented to explain the relationship between BEEP’s predictions in terms of its features for a better understanding of feature importance. The model interpretability starts from the pre-trained BEEP model. The baseline inputs for IG are 4 zero matrices with the same shape to CNN model inputs. TensorFlow’s automatic differentiation capabilities are used to compute the gradients and integrate the gradients along the path from the baseline to the input using numerical integration methods. Once integrated gradients are computed, attribute feature importance by aggregating the integrated gradients across all input features. The feature importance is then visualized by logomarker package.

### Statistical analysis

To proceed with statistical analysis, measurements were generally performed on biological replicates with the number of replicates stated in Methods or Figure legends. Bar plots and associated statistical analyses were generated using Prism 9 (GraphPad). The replicate data were shown as Mean ± SD. Statistical significance was determined using unpaired two-sided *t* test or Fisher’s exact test. Asterisks indicate \*\**p* < 0.01 and \*\*\**p* < 0.001.

## Data availability

The data supporting the results in this study are available within the paper and associated Supplementary Information. The raw and analysed datasets generated in this study are available from the NCBI Gene Expression Omnibus database.

## Code availability

The BEEP website can be accessed at http://beep.weililab.org and the standalone program is available on BitBucket https://bitbucket.org/weililab/beep/src/main/.

## Acknowledgements

This work was supported by the National Natural Science Foundation of China (32071441, 32470673), Guangdong Basic and Applied Basic Research Foundation (2023A1515140084), the 111 Project (B16009) and the Construction Project of Liaoning Provincial Key Laboratory, China (2022JH13/10200026) to T.F., and the research grant from the National Institutes of Health (R01HG010753) to W.L.

## Author contributions

T.F. and W.L. conceived the study. T.F., W.L., X.W. and X.C. designed the research. X.W. performed most of the experiments with the help of H.Z. and Z.C. X.C., Zexu Li and S.M. conducted the bioinformatics analysis. All the authors analyzed the data. T.F., W.L. X.W. and X.C. wrote the manuscript with the input from all the other authors. T.F. and W.L. supervised the study.

---

**Extended Data Fig. 1.**
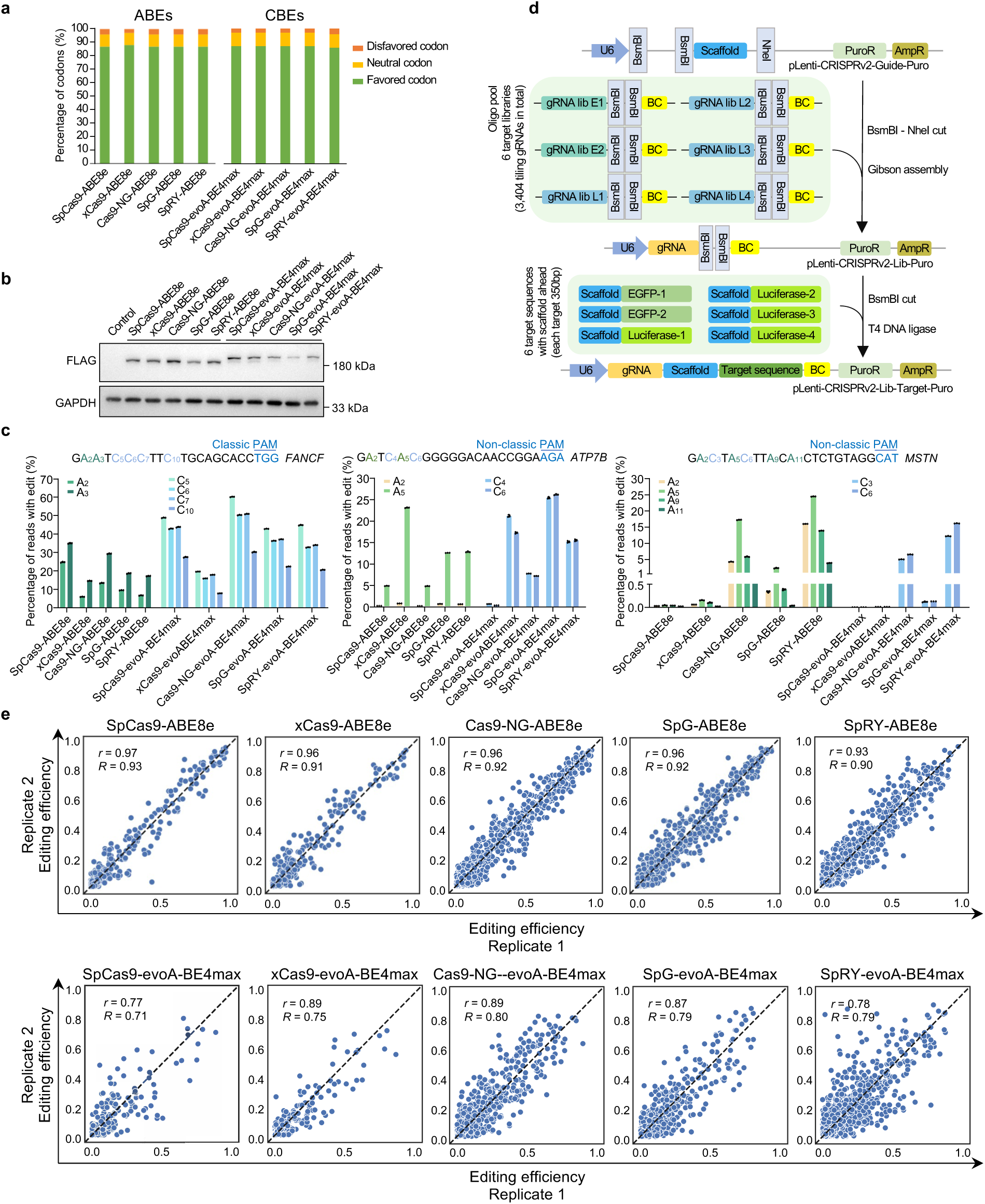
Construction, validation and high-throughput evaluation of a set of BEs. **a**, Percentage of favored, disfavored, and neutral codons across different BE sequences after human codon optimization using GeneScript’s algorithm. **b**, Western blot analysis for A375 cells lentivirally transduced with indicated FLAG-tagged ABEs or CBEs at a low MOI (∼0.1). ‘Control’ sample represents normal un-transduced A375 cells. GAPDH serves as a loading control. **c**, The editing (A>G or C>T) frequency (%) in indicated ABE- or CBE-expressing A375 cells lentivirally transduced with indivudial gRNAs targeting endogenous *FANCF, ATP7B* and *MSTN* genomic sites. Mean ± SD with n = 3 biological replicates. **d**, Schematic representation of the two-step strategy for constructing the BE evaluation library. **e**, Correlation of base editing activities on targets with default PAM patterns for indicated BEs between the two biological replicates. *R* means Spearman correlation coefficient and *r* is the Pearson correlation coefficient.

**Extended Data Fig. 2.**
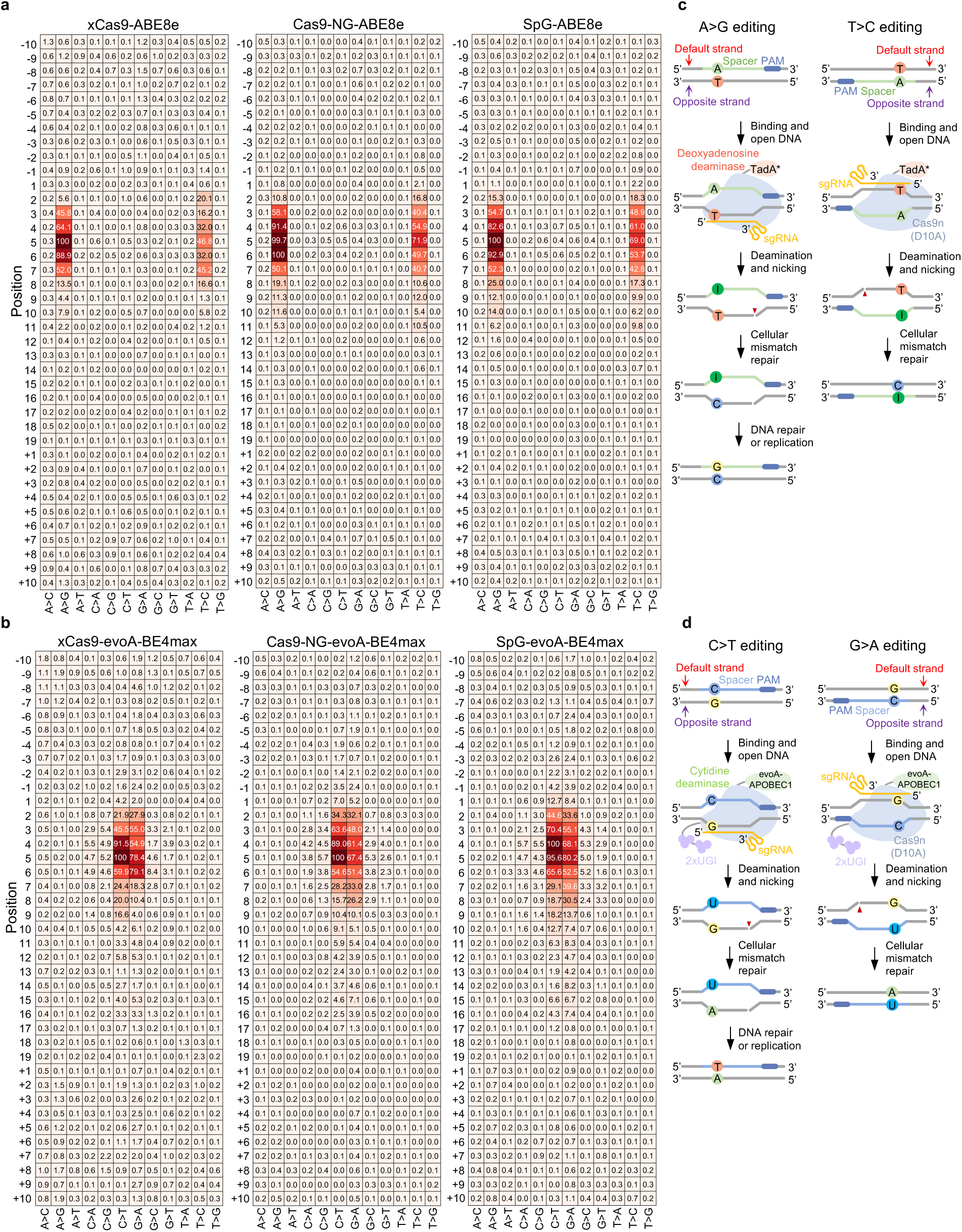
Base editing activity profiles for indicated BEs. **a**,**b**, Heatmap showing the base editing window for indicated ABEs (**a**) and CBEs (**b**). A>G/C>T indicates the spacer (gRNA) and target aligning at the same strand, while T>C/G>A means the spacer and target aligning at different strands. Values in the heatmap indicate the normalized editing efficiency (percentage to the maximal editing). **c**,**d**, Schematic diagram showing the difference between default A>G or C>T editing versus complementary T>C or G>A editing by ABE (**c**) or CBE (**d**), respectively.

**Extended Data Fig. 3.**
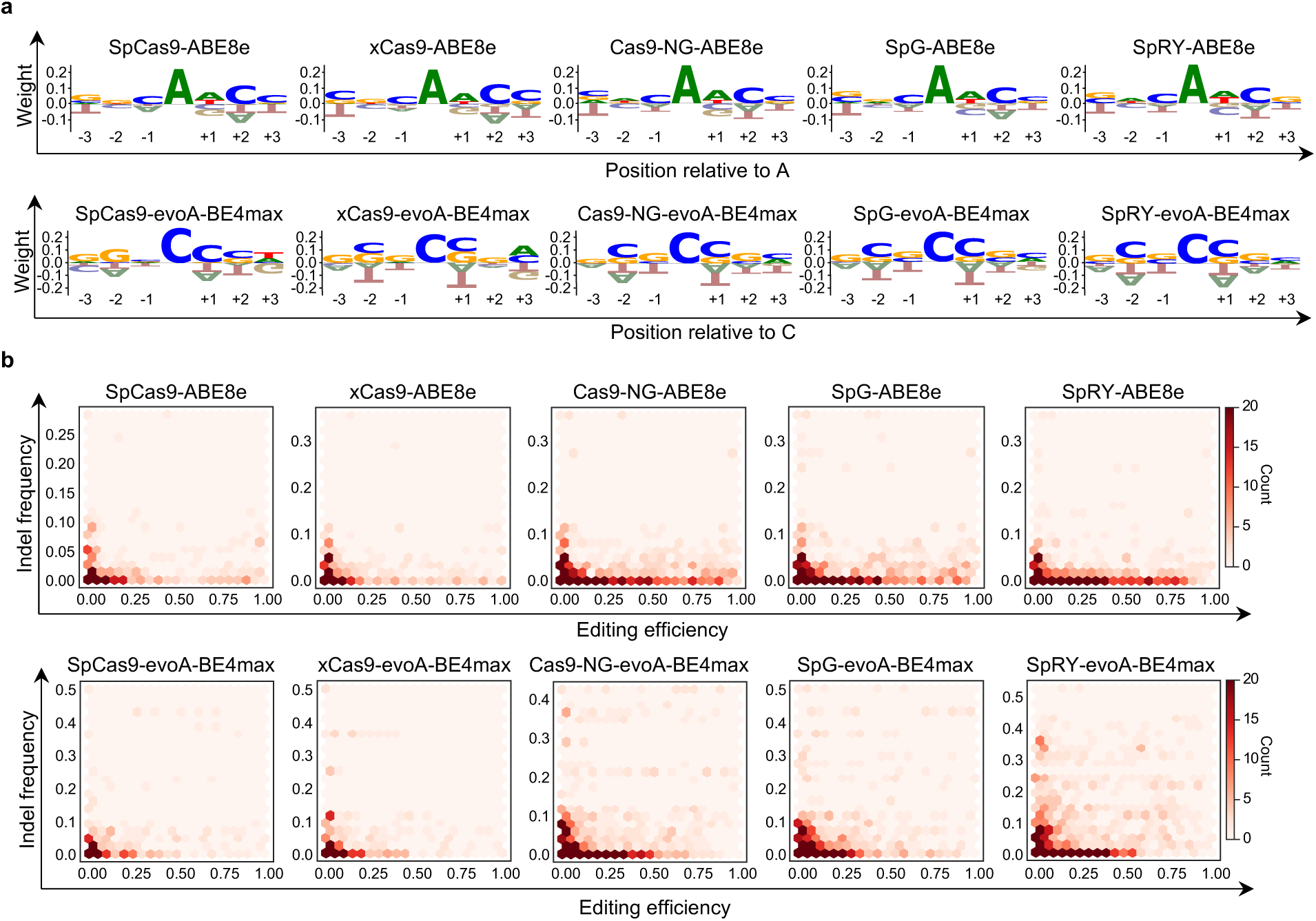
Evaluation of base editing purity and indel for these BEs. **a**, The nearby nucleotide preference of base editing purity. The y-axis is normalized position-specific score. The x-axis is relative position to intended editing nucleotide. The central ‘A’ or ‘C’ is the target position. **b**, Relationship between base editing efficiency and indel frequency for each BE. The x-axis is base editing efficiency and y-axis is the indel frequency. The color legend indicates the number of base editing events.

**Extended Data Fig. 4.**
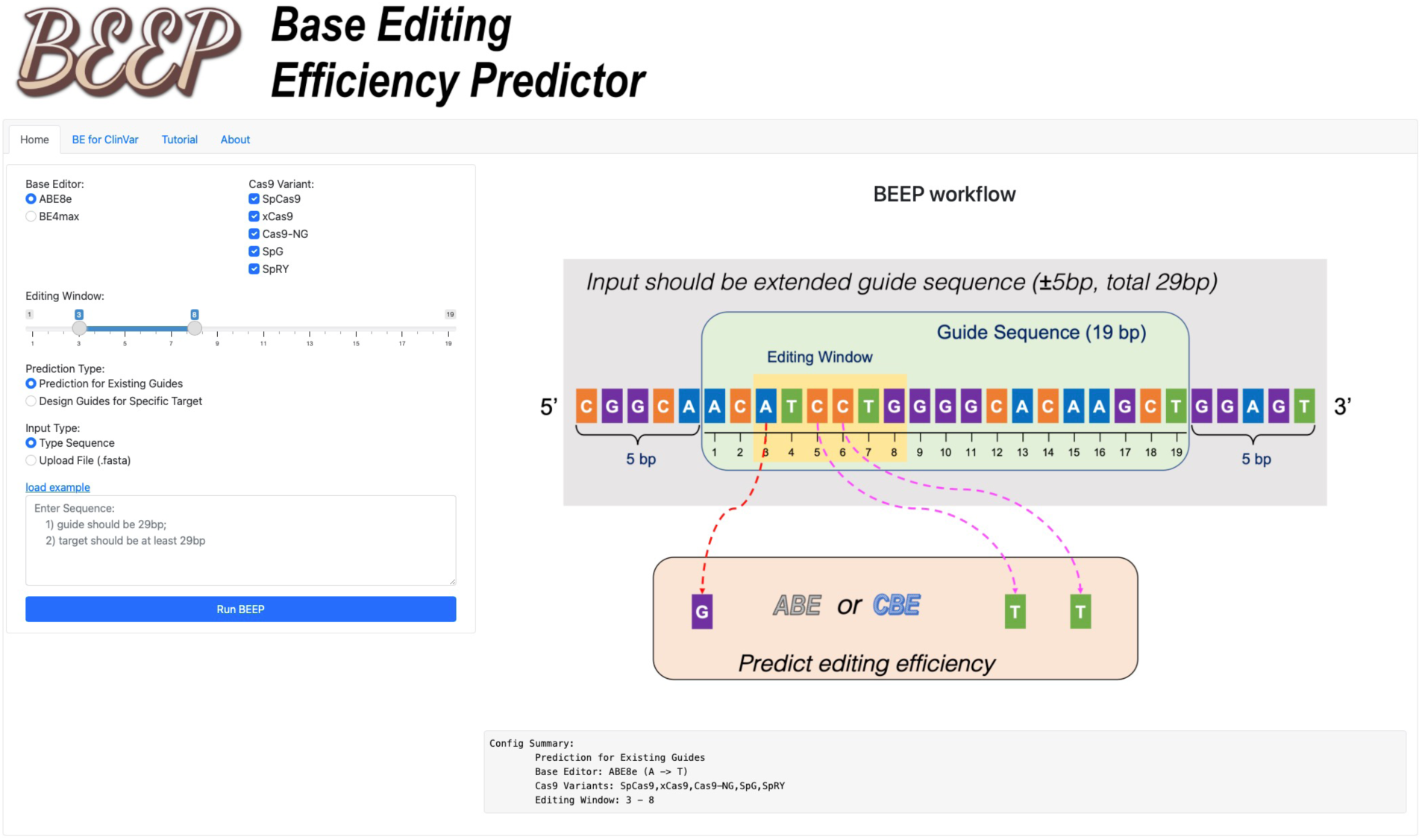
A BEEP-integrated web server for freely public access. Snapshot of a representative page on BEEP web server at http://beep.weililab.org/.

**Extended Data Fig. 5.**
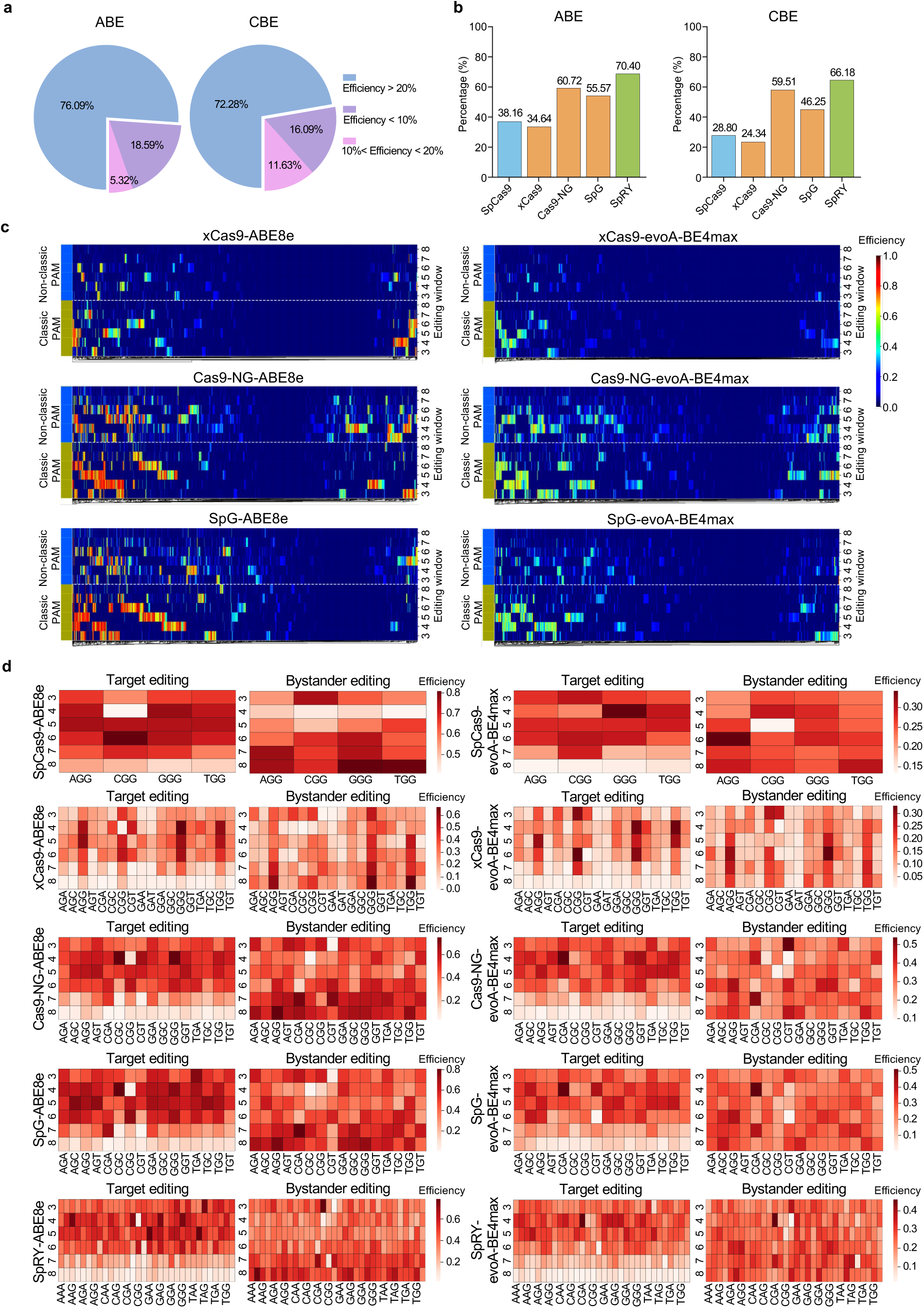
Non-canonical PAM and bystander base editing during high-throughput ClinVar SNV installment. **a**, Pie chart showing the percentage of successfully installed ClinVar SNVs by our ABEs/CBEs in total. **b**, The percentage of successfully installed SNVs with an editing efficiency above 0.2 by indicated ABEs/CBEs. **c**, Heatmap showing the base editing performance of indicated BEs for targets with classic and non-classic PAMs in ClinVar SNV modeling data. **d**, Heatmap showing the relationship between target or bystander editing efficiency and different PAM patterns for indicated BEs using ClinVar SNV modeling data.

**Extended Data Fig. 6.**
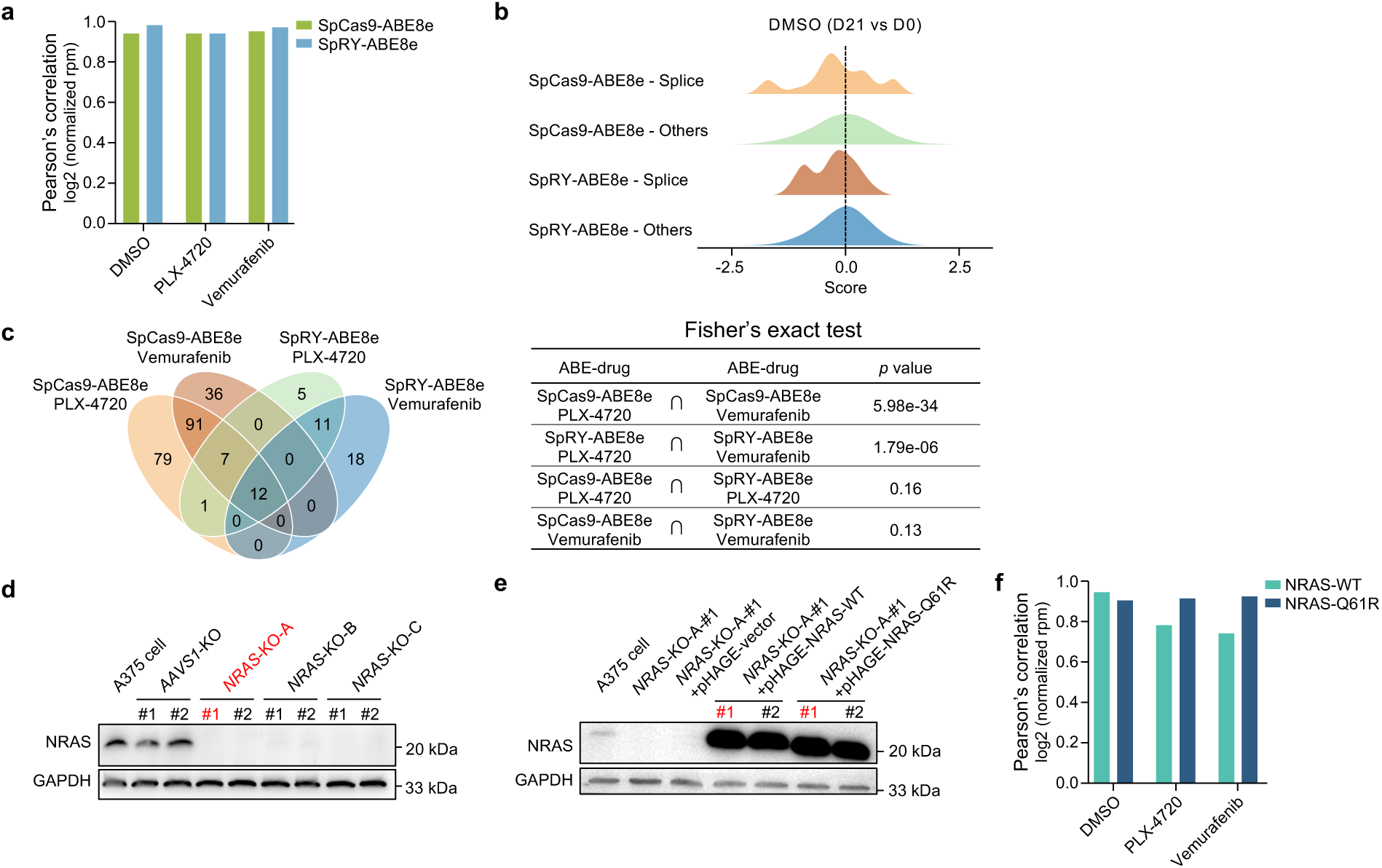
ABE drug resistance screens and druggable gene knockout screens. **a**, Pearson’s correlation for indicated samples between the two biological replicates of ABE drug resistance screens. **b**, Density plot of β-score for gRNAs targeting splice sites of top essential genes in A375 cells or other gRNAs in the library of ABE screening samples (Day 21 vs Day 0) with Vehicle DMSO treatment. **c**, Venn diagram showing the overlap of drug-resistant variant hits across four drug resistance BE screens with two ABEs (SpCas9-ABE8, SpRY-ABE8e) and two drugs (PLX-4720, Vemurafenib). The significance of overlap between indicated groups was shown on the right with Fisher’s exact test *p* values. **d**, Western blot analysis of NRAS expression in indicated single clonal cells to confirm the knockout effect of *NRAS*. Three different gRNAs (A, B and C) targeting *NRAS* under pLenti-CRISPRv2-Puro vector were used to knockout endogenous *NRAS* in A375 cells and two independent single clones (#1 and #2) were expanded for each gRNA. GAPDH serves as a loading control. The red-colored sample was employed for further use. **e**, Western blot analysis of NRAS expression in indicated samples. Vector control, PAM-mutated NRAS-Q61R or NRAS-WT was re-introducted into *NRAS* knockout single clonal cells (*NRAS*-KO-A #1) and two independent single clones (#1 and #2) with replenished NRAS were analyzed. The red colored clones were used in Figs. 6f-i. **f**, Pearson’s correlation for indicated samples between the two biological replicates in druggable gene knockout screens.

