## Supplementary Note 1 for "A comprehensive benchmark for multiple highly efficient base editors with broad targeting scope"

**Supplementary Note 1.** DNA sequences of core components of base editor plasmids used in this study. The base editor-encoding fragments are human codon-optimized and directly synthesized. Sequences for **a**, SpCas9-ABE8e. **b**, xCas9-ABE8e. **c**, Cas9-NG-ABE8e. **d**, SpG-ABE8e. **e**, SpRY-ABE8e. **f**, SpCas9-evoA-BE4max. **g**, xCas9-evoA-BE4max. **h**, Cas9-NG-evoA-BE4max. **i**, SpG-evoA-BE4max. **j**, SpRY-evoA-BE4max.

EF-1α core promoter

SV40 NLS

Cas9 variants (mutation nucleotides in capital letter on top of SpCas9)

TadA-8e

evoAPOBEC1

UGI

Nucleoplasmin NLS

FLAG

P2A

BlastR (blasticidin selection marker)

**a**, pLenti-SpCas9-ABE8e-Blast

gggcagagcgcacatcgcccacagtccccgagaagttggggggaggggtcggcaattgatccggtgcctagagaaggtggcgcggggtaaactgggaaagtgatgtcgtgtactggctccgcctttttcccgagggtgggggagaaccgtatataagtgcagtagtcgccgtgaacgttctttttcgcaacgggtttgccgccagaacacaggaccggttctagatgaaacggacagccgacggaagcgagttcgagtcaccaaagaagaagcggaaagtctctgaggtggagttttcccacgagtactggatgagacatgccctgaccctggccaagagggcacgggatgagagggaggtgcctgtgggagccgtgctggtgctgaacaatagagtgatcggcgagggctggaacagagccatcggcctgcacgacccaacagcccatgccgaaattatggccctgagacagggcggcctggtcatgcagaactacagactgattgacgccaccctgtacgtgacattcgagccttgcgtgatgtgcgccggcgccatgatccactctaggatcggccgcgtggtgtttggcgtgaggaactcaaaaagaggcgccgcaggctccctgatgaacgtgctgaactaccccggcatgaatcaccgcgtcgaaattaccgagggaatcctggcagatgaatgtgccgccctgctgtgcgatttctatcggatgcctagacaggtgttcaatgctcagaagaaggcccagagctccatcaactccggaggatctagcggaggctcctctggctctgagacacctggcacaagcgagagcgcaacacctgaaagcagcgggggcagcagcggggggtcagacaagaagtacagcatcggcctggccatcggcaccaactctgtgggctgggccgtgatcaccgacgagtacaaggtgcccagcaagaaattcaaggtgctgggcaacaccgaccggcacagcatcaagaagaacctgatcggagccctgctgttcgacagcggcgaaacagccgaggccacccggctgaagagaaccgccagaagaagatacaccagacggaagaaccggatctgctatctgcaagagatcttcagcaacgagatggccaaggtggacgacagcttcttccacagactggaagagtccttcctggtggaagaggataagaagcacgagcggcaccccatcttcggcaacatcgtggacgaggtggcctaccacgagaagtaccccaccatctaccacctgagaaagaaactggtggacagcaccgacaaggccgacctgcggctgatctatctggccctggcccacatgatcaagttccggggccacttcctgatcgagggcgacctgaaccccgacaacagcgacgtggacaagctgttcatccagctggtgcagacctacaaccagctgttcgaggaaaaccccatcaacgccagcggcgtggacgccaaggccatcctgtctgccagactgagcaagagcagacggctggaaaatctgatcgcccagctgcccggcgagaagaagaatggcctgttcggaaacctgattgccctgagcctgggcctgacccccaacttcaagagcaacttcgacctggccgaggatgccaaactgcagctgagcaaggacacctacgacgacgacctggacaacctgctggcccagatcggcgaccagtacgccgacctgtttctggccgccaagaacctgtccgacgccatcctgctgagcgacatcctgagagtgaacaccgagatcaccaaggcccccctgagcgcctctatgatcaagagatacgacgagcaccaccaggacctgaccctgctgaaagctctcgtgcggcagcagctgcctgagaagtacaaagagattttcttcgaccagagcaagaacggctacgccggctacattgacggcggagccagccaggaagagttctacaagttcatcaagcccatcctggaaaagatggacggcaccgaggaactgctcgtgaagctgaacagagaggacctgctgcggaagcagcggaccttcgacaacggcagcatcccccaccagatccacctgggagagctgcacgccattctgcggcggcaggaagatttttacccattcctgaaggacaaccgggaaaagatcgagaagatcctgaccttccgcatcccctactacgtgggccctctggccaggggaaacagcagattcgcctggatgaccagaaagagcgaggaaaccatcaccccctggaacttcgaggaagtggtggacaagggcgcttccgcccagagcttcatcgagcggatgaccaacttcgataagaacctgcccaacgagaaggtgctgcccaagcacagcctgctgtacgagtacttcaccgtgtataacgagctgaccaaagtgaaatacgtgaccgagggaatgagaaagcccgccttcctgagcggcgagcagaaaaaggccatcgtggacctgctgttcaagaccaaccggaaagtgaccgtgaagcagctgaaagaggactacttcaagaaaatcgagtgcttcgactccgtggaaatctccggcgtggaagatcggttcaacgcctccctgggcacataccacgatctgctgaaaattatcaaggacaaggacttcctggacaatgaggaaaacgaggacattctggaagatatcgtgctgaccctgacactgtttgaggacagagagatgatcgaggaacggctgaaaacctatgcccacctgttcgacgacaaagtgatgaagcagctgaagcggcggagatacaccggctggggcaggctgagccggaagctgatcaacggcatccgggacaagcagtccggcaagacaatcctggatttcctgaagtccgacggcttcgccaacagaaacttcatgcagctgatccacgacgacagcctgacctttaaagaggacatccagaaagcccaggtgtccggccagggcgatagcctgcacgagcacattgccaatctggccggcagccccgccattaagaagggcatcctgcagacagtgaaggtggtggacgagctcgtgaaagtgatgggccggcacaagcccgagaacatcgtgatcgaaatggccagagagaaccagaccacccagaagggacagaagaacagccgcgagagaatgaagcggatcgaagagggcatcaaagagctgggcagccagatcctgaaagaacaccccgtggaaaacacccagctgcagaacgagaagctgtacctgtactacctgcagaatgggcgggatatgtacgtggaccaggaactggacatcaaccggctgtccgactacgatgtggaccatatcgtgcctcagagctttctgaaggacgactccatcgacaacaaggtgctgaccagaagcgacaagaaccggggcaagagcgacaacgtgccctccgaagaggtcgtgaagaagatgaagaactactggcggcagctgctgaacgccaagctgattacccagagaaagttcgacaatctgaccaaggccgagagaggcggcctgagcgaactggataaggccggcttcatcaagagacagctggtggaaacccggcagatcacaaagcacgtggcacagatcctggactcccggatgaacactaagtacgacgagaatgacaagctgatccgggaagtgaaagtgatcaccctgaagtccaagctggtgtccgatttccggaaggatttccagttttacaaagtgcgcgagatcaacaactaccaccacgcccacgacgcctacctgaacgccgtcgtgggaaccgccctgatcaaaaagtaccctaagctggaaagcgagttcgtgtacggcgactacaaggtgtacgacgtgcggaagatgatcgccaagagcgagcaggaaatcggcaaggctaccgccaagtacttcttctacagcaacatcatgaactttttcaagaccgagattaccctggccaacggcgagatccggaagcggcctctgatcgagacaaacggcgaaaccggggagatcgtgtgggataagggccgggattttgccaccgtgcggaaagtgctgagcatgccccaagtgaatatcgtgaaaaagaccgaggtgcagacaggcggcttcagcaaagagtctatcctgcccaagaggaacagcgataagctgatcgccagaaagaaggactgggaccctaagaagtacggcggcttcgacagccccaccgtggcctattctgtgctggtggtggccaaagtggaaaagggcaagtccaagaaactgaagagtgtgaaagagctgctggggatcaccatcatggaaagaagcagcttcgagaagaatcccatcgactttctggaagccaagggctacaaagaagtgaaaaaggacctgatcatcaagctgcctaagtactccctgttcgagctggaaaacggccggaagagaatgctggcctctgccggcgaactgcagaagggaaacgaactggccctgccctccaaatatgtgaacttcctgtacctggccagccactatgagaagctgaagggctcccccgaggataatgagcagaaacagctgtttgtggaacagcacaagcactacctggacgagatcatcgagcagatcagcgagttctccaagagagtgatcctggccgacgctaatctggacaaagtgctgtccgcctacaacaagcaccgggataagcccatcagagagcaggccgagaatatcatccacctgtttaccctgaccaatctgggagcccctgccgccttcaagtactttgacaccaccatcgaccggaagaggtacaccagcaccaaagaggtgctggacgccaccctgatccaccagagcatcaccggcctgtacgagacacggatcgacctgtctcagctgggaggcgactctggtggttctaagcgacctgccgccacaaagaaggctggacaggctaagaagaagaaagattacaaagacgatgacgataagggatccggcgcaacaaacttctctctgctgaaacaagccggagatgtcgaagagaatcctggaccgatggccaagcctttgtctcaagaagaatccaccctcattgaaagagcaacggctacaatcaacagcatccccatctctgaagactacagcgtcgccagcgcagctctctctagcgacggccgcatcttcactggtgtcaatgtatatcattttactgggggaccttgtgcagaactcgtggtgctgggcactgctgctgctgcggcagctggcaacctgacttgtatcgtcgcgatcggaaatgagaacaggggcatcttgagcccctgcggacggtgccgacaggtgcttctcgatctgcatcctgggatcaaagccatagtgaaggacagtgatggacagccgacggcagttgggattcgtgaattgctgccctctggttatgtgtgggagggctaa

**b**, pLenti-xCas9-ABE8e-Blast

gggcagagcgcacatcgcccacagtccccgagaagttggggggaggggtcggcaattgatccggtgcctagagaaggtggcgcggggtaaactgggaaagtgatgtcgtgtactggctccgcctttttcccgagggtgggggagaaccgtatataagtgcagtagtcgccgtgaacgttctttttcgcaacgggtttgccgccagaacacaggaccggttctagatgaaacggacagccgacggaagcgagttcgagtcaccaaagaagaagcggaaagtctctgaggtggagttttcccacgagtactggatgagacatgccctgaccctggccaagagggcacgggatgagagggaggtgcctgtgggagccgtgctggtgctgaacaatagagtgatcggcgagggctggaacagagccatcggcctgcacgacccaacagcccatgccgaaattatggccctgagacagggcggcctggtcatgcagaactacagactgattgacgccaccctgtacgtgacattcgagccttgcgtgatgtgcgccggcgccatgatccactctaggatcggccgcgtggtgtttggcgtgaggaactcaaaaagaggcgccgcaggctccctgatgaacgtgctgaactaccccggcatgaatcaccgcgtcgaaattaccgagggaatcctggcagatgaatgtgccgccctgctgtgcgatttctatcggatgcctagacaggtgttcaatgctcagaagaaggcccagagctccatcaactccggaggatctagcggaggctcctctggctctgagacacctggcacaagcgagagcgcaacacctgaaagcagcgggggcagcagcggggggtcagacaagaagtacagcatcggcctggCcatcggcaccaactctgtgggctgggccgtgatcaccgacgagtacaaggtgcccagcaagaaattcaaggtgctgggcaacaccgaccggcacagcatcaagaagaacctgatcggagccctgctgttcgacagcggcgaaacagccgaggccacccggctgaagagaaccgccagaagaagatacaccagacggaagaaccggatctgctatctgcaagagatcttcagcaacgagatggccaaggtggacgacagcttcttccacagactggaagagtccttcctggtggaagaggataagaagcacgagcggcaccccatcttcggcaacatcgtggacgaggtggcctaccacgagaagtaccccaccatctaccacctgagaaagaaactggtggacagcaccgacaaggccgacctgcggctgatctatctggccctggcccacatgatcaagttccggggccacttcctgatcgagggcgacctgaaccccgacaacagcgacgtggacaagctgttcatccagctggtgcagacctacaaccagctgttcgaggaaaaccccatcaacgccagcggcgtggacgccaaggccatcctgtctgccagactgagcaagagcagacggctggaaaatctgatcgcccagctgcccggcgagaagaagaatggcctgttcggaaacctgattgccctgagcctgggcctgacccccaacttcaagagcaacttcgacctggccgaggatAccaaactgcagctgagcaaggacacctacgacgacgacctggacaacctgctggcccagatcggcgaccagtacgccgacctgtttctggccgccaagaacctgtccgacgccatcctgctgagcgacatcctgagagtgaacaccgagatcaccaaggcccccctgagcgcctctatgatcaagCTatacgacgagcaccaccaggacctgaccctgctgaaagctctcgtgcggcagcagctgcctgagaagtacaaagagattttcttcgaccagagcaagaacggctacgccggctacattgacggcggagccagccaggaagagttctacaagttcatcaagcccatcctggaaaagatggacggcaccgaggaactgctcgtgaagctgaacagagaggacctgctgcggaagcagcggaccttcgacaacggcaTcatcccccaccagatccacctgggagagctgcacgccattctgcggcggcaggaagatttttacccattcctgaaggacaaccgggaaaagatcgagaagatcctgaccttccgcatcccctactacgtgggccctctggccaggggaaacagcagattcgcctggatgaccagaaagagcgaggaaaccatcaccccctggaacttcgagAaagtggtggacaagggcgcttccgcccagagcttcatcgagcggatgaccaacttcgataagaacctgcccaacgagaaggtgctgcccaagcacagcctgctgtacgagtacttcaccgtgtataacgagctgaccaaagtgaaatacgtgaccgagggaatgagaaagcccgccttcctgagcggcgaTcagaaaaaggccatcgtggacctgctgttcaagaccaaccggaaagtgaccgtgaagcagctgaaagaggactacttcaagaaaatcgagtgcttcgactccgtggaaatctccggcgtggaagatcggttcaacgcctccctgggcacataccacgatctgctgaaaattatcaaggacaaggacttcctggacaatgaggaaaacgaggacattctggaagatatcgtgctgaccctgacactgtttgaggacagagagatgatcgaggaacggctgaaaacctatgcccacctgttcgacgacaaagtgatgaagcagctgaagcggcggagatacaccggctggggcaggctgagccggaagctgatcaacggcatccgggacaagcagtccggcaagacaatcctggatttcctgaagtccgacggcttcgccaacagaaacttcatTcagctgatccacgacgacagcctgacctttaaagaggacatccagaaagcccaggtgtccggccagggcgatagcctgcacgagcacattgccaatctggccggcagccccgccattaagaagggcatcctgcagacagtgaaggtggtggacgagctcgtgaaagtgatgggccggcacaagcccgagaacatcgtgatcgaaatggccagagagaaccagaccacccagaagggacagaagaacagccgcgagagaatgaagcggatcgaagagggcatcaaagagctgggcagccagatcctgaaagaacaccccgtggaaaacacccagctgcagaacgagaagctgtacctgtactacctgcagaatgggcgggatatgtacgtggaccaggaactggacatcaaccggctgtccgactacgatgtggaccatatcgtgcctcagagctttctgaaggacgactccatcgacaacaaggtgctgaccagaagcgacaagaaccggggcaagagcgacaacgtgccctccgaagaggtcgtgaagaagatgaagaactactggcggcagctgctgaacgccaagctgattacccagagaaagttcgacaatctgaccaaggccgagagaggcggcctgagcgaactggataaggccggcttcatcaagagacagctggtggaaacccggcagatcacaaagcacgtggcacagatcctggactcccggatgaacactaagtacgacgagaatgacaagctgatccgggaagtgaaagtgatcaccctgaagtccaagctggtgtccgatttccggaaggatttccagttttacaaagtgcgcgagatcaacaactaccaccacgcccacgacgcctacctgaacgccgtcgtgggaaccgccctgatcaaaaagtaccctaagctggaaagcgagttcgtgtacggcgactacaaggtgtacgacgtgcggaagatgatcgccaagagcgagcaggaaatcggcaaggctaccgccaagtacttcttctacagcaacatcatgaactttttcaagaccgagattaccctggccaacggcgagatccggaagcggcctctgatcgagacaaacggcgaaaccggggagatcgtgtgggataagggccgggattttgccaccgtgcggaaagtgctgagcatgccccaagtgaatatcgtgaaaaagaccgaggtgcagacaggcggcttcagcaaagagtctatcctgcccaagaggaacagcgataagctgatcgccagaaagaaggactgggaccctaagaagtacggcggcttcgacagccccaccgtggcctattctgtgctggtggtggccaaagtggaaaagggcaagtccaagaaactgaagagtgtgaaagagctgctggggatcaccatcatggaaagaagcagcttcgagaagaatcccatcgactttctggaagccaagggctacaaagaagtgaaaaaggacctgatcatcaagctgcctaagtactccctgttcgagctggaaaacggccggaagagaatgctggcctctgccggcgTactgcagaagggaaacgaactggccctgccctccaaatatgtgaacttcctgtacctggccagccactatgagaagctgaagggctcccccgaggataatgagcagaaacagctgtttgtggaacagcacaagcactacctggacgagatcatcgagcagatcagcgagttctccaagagagtgatcctggccgacgctaatctggacaaagtgctgtccgcctacaacaagcaccgggataagcccatcagagagcaggccgagaatatcatccacctgtttaccctgaccaatctgggagcccctgccgccttcaagtactttgacaccaccatcgaccggaagaggtacaccagcaccaaagaggtgctggacgccaccctgatccaccagagcatcaccggcctgtacgagacacggatcgacctgtctcagctgggaggcgactctggtggttctaagcgacctgccgccacaaagaaggctggacaggctaagaagaagaaagattacaaagacgatgacgataagggatccggcgcaacaaacttctctctgctgaaacaagccggagatgtcgaagagaatcctggaccgatggccaagcctttgtctcaagaagaatccaccctcattgaaagagcaacggctacaatcaacagcatccccatctctgaagactacagcgtcgccagcgcagctctctctagcgacggccgcatcttcactggtgtcaatgtatatcattttactgggggaccttgtgcagaactcgtggtgctgggcactgctgctgctgcggcagctggcaacctgacttgtatcgtcgcgatcggaaatgagaacaggggcatcttgagcccctgcggacggtgccgacaggtgcttctcgatctgcatcctgggatcaaagccatagtgaaggacagtgatggacagccgacggcagttgggattcgtgaattgctgccctctggttatgtgtgggagggctaa

**c**, pLenti-Cas9-NG-ABE8e-Blast

gggcagagcgcacatcgcccacagtccccgagaagttggggggaggggtcggcaattgatccggtgcctagagaaggtggcgcggggtaaactgggaaagtgatgtcgtgtactggctccgcctttttcccgagggtgggggagaaccgtatataagtgcagtagtcgccgtgaacgttctttttcgcaacgggtttgccgccagaacacaggaccggttctagatgaaacggacagccgacggaagcgagttcgagtcaccaaagaagaagcggaaagtctctgaggtggagttttcccacgagtactggatgagacatgccctgaccctggccaagagggcacgggatgagagggaggtgcctgtgggagccgtgctggtgctgaacaatagagtgatcggcgagggctggaacagagccatcggcctgcacgacccaacagcccatgccgaaattatggccctgagacagggcggcctggtcatgcagaactacagactgattgacgccaccctgtacgtgacattcgagccttgcgtgatgtgcgccggcgccatgatccactctaggatcggccgcgtggtgtttggcgtgaggaactcaaaaagaggcgccgcaggctccctgatgaacgtgctgaactaccccggcatgaatcaccgcgtcgaaattaccgagggaatcctggcagatgaatgtgccgccctgctgtgcgatttctatcggatgcctagacaggtgttcaatgctcagaagaaggcccagagctccatcaactccggaggatctagcggaggctcctctggctctgagacacctggcacaagcgagagcgcaacacctgaaagcagcgggggcagcagcggggggtcagacaagaagtacagcatcggcctggccatcggcaccaactctgtgggctgggccgtgatcaccgacgagtacaaggtgcccagcaagaaattcaaggtgctgggcaacaccgaccggcacagcatcaagaagaacctgatcggagccctgctgttcgacagcggcgaaacagccgaggccacccggctgaagagaaccgccagaagaagatacaccagacggaagaaccggatctgctatctgcaagagatcttcagcaacgagatggccaaggtggacgacagcttcttccacagactggaagagtccttcctggtggaagaggataagaagcacgagcggcaccccatcttcggcaacatcgtggacgaggtggcctaccacgagaagtaccccaccatctaccacctgagaaagaaactggtggacagcaccgacaaggccgacctgcggctgatctatctggccctggcccacatgatcaagttccggggccacttcctgatcgagggcgacctgaaccccgacaacagcgacgtggacaagctgttcatccagctggtgcagacctacaaccagctgttcgaggaaaaccccatcaacgccagcggcgtggacgccaaggccatcctgtctgccagactgagcaagagcagacggctggaaaatctgatcgcccagctgcccggcgagaagaagaatggcctgttcggaaacctgattgccctgagcctgggcctgacccccaacttcaagagcaacttcgacctggccgaggatgccaaactgcagctgagcaaggacacctacgacgacgacctggacaacctgctggcccagatcggcgaccagtacgccgacctgtttctggccgccaagaacctgtccgacgccatcctgctgagcgacatcctgagagtgaacaccgagatcaccaaggcccccctgagcgcctctatgatcaagagatacgacgagcaccaccaggacctgaccctgctgaaagctctcgtgcggcagcagctgcctgagaagtacaaagagattttcttcgaccagagcaagaacggctacgccggctacattgacggcggagccagccaggaagagttctacaagttcatcaagcccatcctggaaaagatggacggcaccgaggaactgctcgtgaagctgaacagagaggacctgctgcggaagcagcggaccttcgacaacggcagcatcccccaccagatccacctgggagagctgcacgccattctgcggcggcaggaagatttttacccattcctgaaggacaaccgggaaaagatcgagaagatcctgaccttccgcatcccctactacgtgggccctctggccaggggaaacagcagattcgcctggatgaccagaaagagcgaggaaaccatcaccccctggaacttcgaggaagtggtggacaagggcgcttccgcccagagcttcatcgagcggatgaccaacttcgataagaacctgcccaacgagaaggtgctgcccaagcacagcctgctgtacgagtacttcaccgtgtataacgagctgaccaaagtgaaatacgtgaccgagggaatgagaaagcccgccttcctgagcggcgagcagaaaaaggccatcgtggacctgctgttcaagaccaaccggaaagtgaccgtgaagcagctgaaagaggactacttcaagaaaatcgagtgcttcgactccgtggaaatctccggcgtggaagatcggttcaacgcctccctgggcacataccacgatctgctgaaaattatcaaggacaaggacttcctggacaatgaggaaaacgaggacattctggaagatatcgtgctgaccctgacactgtttgaggacagagagatgatcgaggaacggctgaaaacctatgcccacctgttcgacgacaaagtgatgaagcagctgaagcggcggagatacaccggctggggcaggctgagccggaagctgatcaacggcatccgggacaagcagtccggcaagacaatcctggatttcctgaagtccgacggcttcgccaacagaaacttcatgcagctgatccacgacgacagcctgacctttaaagaggacatccagaaagcccaggtgtccggccagggcgatagcctgcacgagcacattgccaatctggccggcagccccgccattaagaagggcatcctgcagacagtgaaggtggtggacgagctcgtgaaagtgatgggccggcacaagcccgagaacatcgtgatcgaaatggccagagagaaccagaccacccagaagggacagaagaacagccgcgagagaatgaagcggatcgaagagggcatcaaagagctgggcagccagatcctgaaagaacaccccgtggaaaacacccagctgcagaacgagaagctgtacctgtactacctgcagaatgggcgggatatgtacgtggaccaggaactggacatcaaccggctgtccgactacgatgtggaccatatcgtgcctcagagctttctgaaggacgactccatcgacaacaaggtgctgaccagaagcgacaagaaccggggcaagagcgacaacgtgccctccgaagaggtcgtgaagaagatgaagaactactggcggcagctgctgaacgccaagctgattacccagagaaagttcgacaatctgaccaaggccgagagaggcggcctgagcgaactggataaggccggcttcatcaagagacagctggtggaaacccggcagatcacaaagcacgtggcacagatcctggactcccggatgaacactaagtacgacgagaatgacaagctgatccgggaagtgaaagtgatcaccctgaagtccaagctggtgtccgatttccggaaggatttccagttttacaaagtgcgcgagatcaacaactaccaccacgcccacgacgcctacctgaacgccgtcgtgggaaccgccctgatcaaaaagtaccctaagctggaaagcgagttcgtgtacggcgactacaaggtgtacgacgtgcggaagatgatcgccaagagcgagcaggaaatcggcaaggctaccgccaagtacttcttctacagcaacatcatgaactttttcaagaccgagattaccctggccaacggcgagatccggaagcggcctctgatcgagacaaacggcgaaaccggggagatcgtgtgggataagggccgggattttgccaccgtgcggaaagtgctgagcatgccccaagtgaatatcgtgaaaaagaccgaggtgcagacaggcggcttcagcaaagagtctatccggcccaagaggaacagcgataagctgatcgccagaaagaaggactgggaccctaagaagtacggcggcttcgtcagccccaccgtggcctattctgtgctggtggtggccaaagtggaaaagggcaagtccaagaaactgaagagtgtgaaagagctgctggggatcaccatcatggaaagaagcagcttcgagaagaatcccatcgactttctggaagccaagggctacaaagaagtgaaaaaggacctgatcatcaagctgcctaagtactccctgttcgagctggaaaacggccggaagagaatgctggcctctgcccgcttcctgcagaagggaaacgaactggccctgccctccaaatatgtgaacttcctgtacctggccagccactatgagaagctgaagggctcccccgaggataatgagcagaaacagctgtttgtggaacagcacaagcactacctggacgagatcatcgagcagatcagcgagttctccaagagagtgatcctggccgacgctaatctggacaaagtgctgtccgcctacaacaagcaccgggataagcccatcagagagcaggccgagaatatcatccacctgtttaccctgaccaatctgggagcccctcgcgccttcaagtactttgacaccaccatcgaccggaaggtgtacaggagcaccaaagaggtgctggacgccaccctgatccaccagagcatcaccggcctgtacgagacacggatcgacctgtctcagctgggaggcgactctggtggttctaagcgacctgccgccacaaagaaggctggacaggctaagaagaagaaagattacaaagacgatgacgataagggatccggcgcaacaaacttctctctgctgaaacaagccggagatgtcgaagagaatcctggaccgatggccaagcctttgtctcaagaagaatccaccctcattgaaagagcaacggctacaatcaacagcatccccatctctgaagactacagcgtcgccagcgcagctctctctagcgacggccgcatcttcactggtgtcaatgtatatcattttactgggggaccttgtgcagaactcgtggtgctgggcactgctgctgctgcggcagctggcaacctgacttgtatcgtcgcgatcggaaatgagaacaggggcatcttgagcccctgcggacggtgccgacaggtgcttctcgatctgcatcctgggatcaaagccatagtgaaggacagtgatggacagccgacggcagttgggattcgtgaattgctgccctctggttatgtgtgggagggctaa

**d**, pLenti-SpG-ABE8e-Blast

gggcagagcgcacatcgcccacagtccccgagaagttggggggaggggtcggcaattgatccggtgcctagagaaggtggcgcggggtaaactgggaaagtgatgtcgtgtactggctccgcctttttcccgagggtgggggagaaccgtatataagtgcagtagtcgccgtgaacgttctttttcgcaacgggtttgccgccagaacacaggaccggttctagatgaaacggacagccgacggaagcgagttcgagtcaccaaagaagaagcggaaagtctctgaggtggagttttcccacgagtactggatgagacatgccctgaccctggccaagagggcacgggatgagagggaggtgcctgtgggagccgtgctggtgctgaacaatagagtgatcggcgagggctggaacagagccatcggcctgcacgacccaacagcccatgccgaaattatggccctgagacagggcggcctggtcatgcagaactacagactgattgacgccaccctgtacgtgacattcgagccttgcgtgatgtgcgccggcgccatgatccactctaggatcggccgcgtggtgtttggcgtgaggaactcaaaaagaggcgccgcaggctccctgatgaacgtgctgaactaccccggcatgaatcaccgcgtcgaaattaccgagggaatcctggcagatgaatgtgccgccctgctgtgcgatttctatcggatgcctagacaggtgttcaatgctcagaagaaggcccagagctccatcaactccggaggatctagcggaggctcctctggctctgagacacctggcacaagcgagagcgcaacacctgaaagcagcgggggcagcagcggggggtcagacaagaagtacagcatcggcctggccatcggcaccaactctgtgggctgggccgtgatcaccgacgagtacaaggtgcccagcaagaaattcaaggtgctgggcaacaccgaccggcacagcatcaagaagaacctgatcggagccctgctgttcgacagcggcgaaacagccgaggccacccggctgaagagaaccgccagaagaagatacaccagacggaagaaccggatctgctatctgcaagagatcttcagcaacgagatggccaaggtggacgacagcttcttccacagactggaagagtccttcctggtggaagaggataagaagcacgagcggcaccccatcttcggcaacatcgtggacgaggtggcctaccacgagaagtaccccaccatctaccacctgagaaagaaactggtggacagcaccgacaaggccgacctgcggctgatctatctggccctggcccacatgatcaagttccggggccacttcctgatcgagggcgacctgaaccccgacaacagcgacgtggacaagctgttcatccagctggtgcagacctacaaccagctgttcgaggaaaaccccatcaacgccagcggcgtggacgccaaggccatcctgtctgccagactgagcaagagcagacggctggaaaatctgatcgcccagctgcccggcgagaagaagaatggcctgttcggaaacctgattgccctgagcctgggcctgacccccaacttcaagagcaacttcgacctggccgaggatgccaaactgcagctgagcaaggacacctacgacgacgacctggacaacctgctggcccagatcggcgaccagtacgccgacctgtttctggccgccaagaacctgtccgacgccatcctgctgagcgacatcctgagagtgaacaccgagatcaccaaggcccccctgagcgcctctatgatcaagagatacgacgagcaccaccaggacctgaccctgctgaaagctctcgtgcggcagcagctgcctgagaagtacaaagagattttcttcgaccagagcaagaacggctacgccggctacattgacggcggagccagccaggaagagttctacaagttcatcaagcccatcctggaaaagatggacggcaccgaggaactgctcgtgaagctgaacagagaggacctgctgcggaagcagcggaccttcgacaacggcagcatcccccaccagatccacctgggagagctgcacgccattctgcggcggcaggaagatttttacccattcctgaaggacaaccgggaaaagatcgagaagatcctgaccttccgcatcccctactacgtgggccctctggccaggggaaacagcagattcgcctggatgaccagaaagagcgaggaaaccatcaccccctggaacttcgaggaagtggtggacaagggcgcttccgcccagagcttcatcgagcggatgaccaacttcgataagaacctgcccaacgagaaggtgctgcccaagcacagcctgctgtacgagtacttcaccgtgtataacgagctgaccaaagtgaaatacgtgaccgagggaatgagaaagcccgccttcctgagcggcgagcagaaaaaggccatcgtggacctgctgttcaagaccaaccggaaagtgaccgtgaagcagctgaaagaggactacttcaagaaaatcgagtgcttcgactccgtggaaatctccggcgtggaagatcggttcaacgcctccctgggcacataccacgatctgctgaaaattatcaaggacaaggacttcctggacaatgaggaaaacgaggacattctggaagatatcgtgctgaccctgacactgtttgaggacagagagatgatcgaggaacggctgaaaacctatgcccacctgttcgacgacaaagtgatgaagcagctgaagcggcggagatacaccggctggggcaggctgagccggaagctgatcaacggcatccgggacaagcagtccggcaagacaatcctggatttcctgaagtccgacggcttcgccaacagaaacttcatgcagctgatccacgacgacagcctgacctttaaagaggacatccagaaagcccaggtgtccggccagggcgatagcctgcacgagcacattgccaatctggccggcagccccgccattaagaagggcatcctgcagacagtgaaggtggtggacgagctcgtgaaagtgatgggccggcacaagcccgagaacatcgtgatcgaaatggccagagagaaccagaccacccagaagggacagaagaacagccgcgagagaatgaagcggatcgaagagggcatcaaagagctgggcagccagatcctgaaagaacaccccgtggaaaacacccagctgcagaacgagaagctgtacctgtactacctgcagaatgggcgggatatgtacgtggaccaggaactggacatcaaccggctgtccgactacgatgtggaccatatcgtgcctcagagctttctgaaggacgactccatcgacaacaaggtgctgaccagaagcgacaagaaccggggcaagagcgacaacgtgccctccgaagaggtcgtgaagaagatgaagaactactggcggcagctgctgaacgccaagctgattacccagagaaagttcgacaatctgaccaaggccgagagaggcggcctgagcgaactggataaggccggcttcatcaagagacagctggtggaaacccggcagatcacaaagcacgtggcacagatcctggactcccggatgaacactaagtacgacgagaatgacaagctgatccgggaagtgaaagtgatcaccctgaagtccaagctggtgtccgatttccggaaggatttccagttttacaaagtgcgcgagatcaacaactaccaccacgcccacgacgcctacctgaacgccgtcgtgggaaccgccctgatcaaaaagtaccctaagctggaaagcgagttcgtgtacggcgactacaaggtgtacgacgtgcggaagatgatcgccaagagcgagcaggaaatcggcaaggctaccgccaagtacttcttctacagcaacatcatgaactttttcaagaccgagattaccctggccaacggcgagatccggaagcggcctctgatcgagacaaacggcgaaaccggggagatcgtgtgggataagggccgggattttgccaccgtgcggaaagtgctgagcatgccccaagtgaatatcgtgaaaaagaccgaggtgcagacaggcggcttcagcaaagagtctatcctgcccaagaggaacagcgataagctgatcgccagaaagaaggactgggaccctaagaagtacggcggcttcCTGTgGcccaccgtggcctattctgtgctggtggtggccaaagtggaaaagggcaagtccaagaaactgaagagtgtgaaagagctgctggggatcaccatcatggaaagaagcagcttcgagaagaatcccatcgactttctggaagccaagggctacaaagaagtgaaaaaggacctgatcatcaagctgcctaagtactccctgttcgagctggaaaacggccggaagagaatgctggcctctgccAAGCaGctgcagaagggaaacgaactggccctgccctccaaatatgtgaacttcctgtacctggccagccactatgagaagctgaagggctcccccgaggataatgagcagaaacagctgtttgtggaacagcacaagcactacctggacgagatcatcgagcagatcagcgagttctccaagagagtgatcctggccgacgctaatctggacaaagtgctgtccgcctacaacaagcaccgggataagcccatcagagagcaggccgagaatatcatccacctgtttaccctgaccaatctgggagcccctgccgccttcaagtactttgacaccaccatcgaccggaagCAgtacaGAagcaccaaagaggtgctggacgccaccctgatccaccagagcatcaccggcctgtacgagacacggatcgacctgtctcagctgggaggcgactctggtggttctaagcgacctgccgccacaaagaaggctggacaggctaagaagaagaaagattacaaagacgatgacgataagggatccggcgcaacaaacttctctctgctgaaacaagccggagatgtcgaagagaatcctggaccgatggccaagcctttgtctcaagaagaatccaccctcattgaaagagcaacggctacaatcaacagcatccccatctctgaagactacagcgtcgccagcgcagctctctctagcgacggccgcatcttcactggtgtcaatgtatatcattttactgggggaccttgtgcagaactcgtggtgctgggcactgctgctgctgcggcagctggcaacctgacttgtatcgtcgcgatcggaaatgagaacaggggcatcttgagcccctgcggacggtgccgacaggtgcttctcgatctgcatcctgggatcaaagccatagtgaaggacagtgatggacagccgacggcagttgggattcgtgaattgctgccctctggttatgtgtgggagggctaa

**e**, pLenti-SpRY-ABE8e-Blast

gggcagagcgcacatcgcccacagtccccgagaagttggggggaggggtcggcaattgatccggtgcctagagaaggtggcgcggggtaaactgggaaagtgatgtcgtgtactggctccgcctttttcccgagggtgggggagaaccgtatataagtgcagtagtcgccgtgaacgttctttttcgcaacgggtttgccgccagaacacaggaccggttctagatgaaacggacagccgacggaagcgagttcgagtcaccaaagaagaagcggaaagtctctgaggtggagttttcccacgagtactggatgagacatgccctgaccctggccaagagggcacgggatgagagggaggtgcctgtgggagccgtgctggtgctgaacaatagagtgatcggcgagggctggaacagagccatcggcctgcacgacccaacagcccatgccgaaattatggccctgagacagggcggcctggtcatgcagaactacagactgattgacgccaccctgtacgtgacattcgagccttgcgtgatgtgcgccggcgccatgatccactctaggatcggccgcgtggtgtttggcgtgaggaactcaaaaagaggcgccgcaggctccctgatgaacgtgctgaactaccccggcatgaatcaccgcgtcgaaattaccgagggaatcctggcagatgaatgtgccgccctgctgtgcgatttctatcggatgcctagacaggtgttcaatgctcagaagaaggcccagagctccatcaactccggaggatctagcggaggctcctctggctctgagacacctggcacaagcgagagcgcaacacctgaaagcagcgggggcagcagcggggggtcagacaagaagtacagcatcggcctggccatcggcaccaactctgtgggctgggccgtgatcaccgacgagtacaaggtgcccagcaagaaattcaaggtgctgggcaacaccgaccggcacagcatcaagaagaacctgatcggagccctgctgttcgacagcggcgaaacagccgagAGAacccggctgaagagaaccgccagaagaagatacaccagacggaagaaccggatctgctatctgcaagagatcttcagcaacgagatggccaaggtggacgacagcttcttccacagactggaagagtccttcctggtggaagaggataagaagcacgagcggcaccccatcttcggcaacatcgtggacgaggtggcctaccacgagaagtaccccaccatctaccacctgagaaagaaactggtggacagcaccgacaaggccgacctgcggctgatctatctggccctggcccacatgatcaagttccggggccacttcctgatcgagggcgacctgaaccccgacaacagcgacgtggacaagctgttcatccagctggtgcagacctacaaccagctgttcgaggaaaaccccatcaacgccagcggcgtggacgccaaggccatcctgtctgccagactgagcaagagcagacggctggaaaatctgatcgcccagctgcccggcgagaagaagaatggcctgttcggaaacctgattgccctgagcctgggcctgacccccaacttcaagagcaacttcgacctggccgaggatgccaaactgcagctgagcaaggacacctacgacgacgacctggacaacctgctggcccagatcggcgaccagtacgccgacctgtttctggccgccaagaacctgtccgacgccatcctgctgagcgacatcctgagagtgaacaccgagatcaccaaggcccccctgagcgcctctatgatcaagagatacgacgagcaccaccaggacctgaccctgctgaaagctctcgtgcggcagcagctgcctgagaagtacaaagagattttcttcgaccagagcaagaacggctacgccggctacattgacggcggagccagccaggaagagttctacaagttcatcaagcccatcctggaaaagatggacggcaccgaggaactgctcgtgaagctgaacagagaggacctgctgcggaagcagcggaccttcgacaacggcagcatcccccaccagatccacctgggagagctgcacgccattctgcggcggcaggaagatttttacccattcctgaaggacaaccgggaaaagatcgagaagatcctgaccttccgcatcccctactacgtgggccctctggccaggggaaacagcagattcgcctggatgaccagaaagagcgaggaaaccatcaccccctggaacttcgaggaagtggtggacaagggcgcttccgcccagagcttcatcgagcggatgaccaacttcgataagaacctgcccaacgagaaggtgctgcccaagcacagcctgctgtacgagtacttcaccgtgtataacgagctgaccaaagtgaaatacgtgaccgagggaatgagaaagcccgccttcctgagcggcgagcagaaaaaggccatcgtggacctgctgttcaagaccaaccggaaagtgaccgtgaagcagctgaaagaggactacttcaagaaaatcgagtgcttcgactccgtggaaatctccggcgtggaagatcggttcaacgcctccctgggcacataccacgatctgctgaaaattatcaaggacaaggacttcctggacaatgaggaaaacgaggacattctggaagatatcgtgctgaccctgacactgtttgaggacagagagatgatcgaggaacggctgaaaacctatgcccacctgttcgacgacaaagtgatgaagcagctgaagcggcggagatacaccggctggggcaggctgagccggaagctgatcaacggcatccgggacaagcagtccggcaagacaatcctggatttcctgaagtccgacggcttcgccaacagaaacttcatgcagctgatccacgacgacagcctgacctttaaagaggacatccagaaagcccaggtgtccggccagggcgatagcctgcacgagcacattgccaatctggccggcagccccgccattaagaagggcatcctgcagacagtgaaggtggtggacgagctcgtgaaagtgatgggccggcacaagcccgagaacatcgtgatcgaaatggccagagagaaccagaccacccagaagggacagaagaacagccgcgagagaatgaagcggatcgaagagggcatcaaagagctgggcagccagatcctgaaagaacaccccgtggaaaacacccagctgcagaacgagaagctgtacctgtactacctgcagaatgggcgggatatgtacgtggaccaggaactggacatcaaccggctgtccgactacgatgtggaccatatcgtgcctcagagctttctgaaggacgactccatcgacaacaaggtgctgaccagaagcgacaagaaccggggcaagagcgacaacgtgccctccgaagaggtcgtgaagaagatgaagaactactggcggcagctgctgaacgccaagctgattacccagagaaagttcgacaatctgaccaaggccgagagaggcggcctgagcgaactggataaggccggcttcatcaagagacagctggtggaaacccggcagatcacaaagcacgtggcacagatcctggactcccggatgaacactaagtacgacgagaatgacaagctgatccgggaagtgaaagtgatcaccctgaagtccaagctggtgtccgatttccggaaggatttccagttttacaaagtgcgcgagatcaacaactaccaccacgcccacgacgcctacctgaacgccgtcgtgggaaccgccctgatcaaaaagtaccctaagctggaaagcgagttcgtgtacggcgactacaaggtgtacgacgtgcggaagatgatcgccaagagcgagcaggaaatcggcaaggctaccgccaagtacttcttctacagcaacatcatgaactttttcaagaccgagattaccctggccaacggcgagatccggaagcggcctctgatcgagacaaacggcgaaaccggggagatcgtgtgggataagggccgggattttgccaccgtgcggaaagtgctgagcatgccccaagtgaatatcgtgaaaaagaccgaggtgcagacaggcggcttcagcaaagagtctatcAGAcccaagaggaacagcgataagctgatcgccagaaagaaggactgggaccctaagaagtacggcggcttcCTGTgGcccaccgtggcctattctgtgctggtggtggccaaagtggaaaagggcaagtccaagaaactgaagagtgtgaaagagctgctggggatcaccatcatggaaagaagcagcttcgagaagaatcccatcgactttctggaagccaagggctacaaagaagtgaaaaaggacctgatcatcaagctgcctaagtactccctgttcgagctggaaaacggccggaagagaatgctggcctctgccAAGCaGctgcagaagggaaacgaactggccctgccctccaaatatgtgaacttcctgtacctggccagccactatgagaagctgaagggctcccccgaggataatgagcagaaacagctgtttgtggaacagcacaagcactacctggacgagatcatcgagcagatcagcgagttctccaagagagtgatcctggccgacgctaatctggacaaagtgctgtccgcctacaacaagcaccgggataagcccatcagagagcaggccgagaatatcatccacctgtttaccctgaccaGActgggagcccctAGAgccttcaagtactttgacaccaccatcgaccCCaagCAgtacaGAagcaccaaagaggtgctggacgccaccctgatccaccagagcatcaccggcctgtacgagacacggatcgacctgtctcagctgggaggcgactctggtggttctaagcgacctgccgccacaaagaaggctggacaggctaagaagaagaaagattacaaagacgatgacgataagggatccggcgcaacaaacttctctctgctgaaacaagccggagatgtcgaagagaatcctggaccgatggccaagcctttgtctcaagaagaatccaccctcattgaaagagcaacggctacaatcaacagcatccccatctctgaagactacagcgtcgccagcgcagctctctctagcgacggccgcatcttcactggtgtcaatgtatatcattttactgggggaccttgtgcagaactcgtggtgctgggcactgctgctgctgcggcagctggcaacctgacttgtatcgtcgcgatcggaaatgagaacaggggcatcttgagcccctgcggacggtgccgacaggtgcttctcgatctgcatcctgggatcaaagccatagtgaaggacagtgatggacagccgacggcagttgggattcgtgaattgctgccctctggttatgtgtgggagggctaa

**f**, pLenti-SpCas9-evoA-BE4max-Blast

gggcagagcgcacatcgcccacagtccccgagaagttggggggaggggtcggcaattgatccggtgcctagagaaggtggcgcggggtaaactgggaaagtgatgtcgtgtactggctccgcctttttcccgagggtgggggagaaccgtatataagtgcagtagtcgccgtgaacgttctttttcgcaacgggtttgccgccagaacacaggaccggttctagaatgaaacggacagccgacggaagcgagttcgagtcaccaaagaagaagcggaaagtcagttcaaagactgggcctgtcgccgtcgatccaaccctgcgccgccggattgaacctcacgagtttgaagtgttctttgacccccgggagctgagaaaggagacatgcctgctgtacgagatcaactggggaggcaggcactccatctggaggcacacctctcagaacacaaataagcacgtggaggtgaacttcatcgagaagtttaccacagagcggtacttctgccccaataccagatgtagcatcacatggtttctgagctggtccccttgcggagagtgtagcagggccatcaccgagttcctgtccagatacccaaatgtgacactgtttatctacatcgccaggctgtatcacctggcaaacccaaggaataggcagggcctgcgcgatctgatcagctccggcgtgaccatccagatcatgacagagcaggagtccggctactgctggcacaacttcgtgaattattctcctagcaacgagtcccactggcctaggtacccacacctgtgggtgcgcctgtacgtgctggagctgtattgcatcatcctgggcctgcccccttgtctgaatatcctgcggagaaagcagagccagctgacctcctttacaatcgccctgcagtcttgtcactatcagaggctgccaccccacatcctgtgggccacaggcctgaagtctggcggatctagcggaggatcttctggcagcgagacaccaggaacaagcgagtcagcaacaccagagagcagtggcggcagcagcggcggcagcgacaagaagtacagcatcggcctggccatcggcaccaactctgtgggctgggccgtgatcaccgacgagtacaaggtgcccagcaagaaattcaaggtgctgggcaacaccgaccggcacagcatcaagaagaacctgatcggagccctgctgttcgacagcggcgaaacagccgaggccacccggctgaagagaaccgccagaagaagatacaccagacggaagaaccggatctgctatctgcaagagatcttcagcaacgagatggccaaggtggacgacagcttcttccacagactggaagagtccttcctggtggaagaggataagaagcacgagcggcaccccatcttcggcaacatcgtggacgaggtggcctaccacgagaagtaccccaccatctaccacctgagaaagaaactggtggacagcaccgacaaggccgacctgcggctgatctatctggccctggcccacatgatcaagttccggggccacttcctgatcgagggcgacctgaaccccgacaacagcgacgtggacaagctgttcatccagctggtgcagacctacaaccagctgttcgaggaaaaccccatcaacgccagcggcgtggacgccaaggccatcctgtctgccagactgagcaagagcagacggctggaaaatctgatcgcccagctgcccggcgagaagaagaatggcctgttcggaaacctgattgccctgagcctgggcctgacccccaacttcaagagcaacttcgacctggccgaggatgccaaactgcagctgagcaaggacacctacgacgacgacctggacaacctgctggcccagatcggcgaccagtacgccgacctgtttctggccgccaagaacctgtccgacgccatcctgctgagcgacatcctgagagtgaacaccgagatcaccaaggcccccctgagcgcctctatgatcaagagatacgacgagcaccaccaggacctgaccctgctgaaagctctcgtgcggcagcagctgcctgagaagtacaaagagattttcttcgaccagagcaagaacggctacgccggctacattgacggcggagccagccaggaagagttctacaagttcatcaagcccatcctggaaaagatggacggcaccgaggaactgctcgtgaagctgaacagagaggacctgctgcggaagcagcggaccttcgacaacggcagcatcccccaccagatccacctgggagagctgcacgccattctgcggcggcaggaagatttttacccattcctgaaggacaaccgggaaaagatcgagaagatcctgaccttccgcatcccctactacgtgggccctctggccaggggaaacagcagattcgcctggatgaccagaaagagcgaggaaaccatcaccccctggaacttcgaggaagtggtggacaagggcgcttccgcccagagcttcatcgagcggatgaccaacttcgataagaacctgcccaacgagaaggtgctgcccaagcacagcctgctgtacgagtacttcaccgtgtataacgagctgaccaaagtgaaatacgtgaccgagggaatgagaaagcccgccttcctgagcggcgagcagaaaaaggccatcgtggacctgctgttcaagaccaaccggaaagtgaccgtgaagcagctgaaagaggactacttcaagaaaatcgagtgcttcgactccgtggaaatctccggcgtggaagatcggttcaacgcctccctgggcacataccacgatctgctgaaaattatcaaggacaaggacttcctggacaatgaggaaaacgaggacattctggaagatatcgtgctgaccctgacactgtttgaggacagagagatgatcgaggaacggctgaaaacctatgcccacctgttcgacgacaaagtgatgaagcagctgaagcggcggagatacaccggctggggcaggctgagccggaagctgatcaacggcatccgggacaagcagtccggcaagacaatcctggatttcctgaagtccgacggcttcgccaacagaaacttcatgcagctgatccacgacgacagcctgacctttaaagaggacatccagaaagcccaggtgtccggccagggcgatagcctgcacgagcacattgccaatctggccggcagccccgccattaagaagggcatcctgcagacagtgaaggtggtggacgagctcgtgaaagtgatgggccggcacaagcccgagaacatcgtgatcgaaatggccagagagaaccagaccacccagaagggacagaagaacagccgcgagagaatgaagcggatcgaagagggcatcaaagagctgggcagccagatcctgaaagaacaccccgtggaaaacacccagctgcagaacgagaagctgtacctgtactacctgcagaatgggcgggatatgtacgtggaccaggaactggacatcaaccggctgtccgactacgatgtggaccatatcgtgcctcagagctttctgaaggacgactccatcgacaacaaggtgctgaccagaagcgacaagaaccggggcaagagcgacaacgtgccctccgaagaggtcgtgaagaagatgaagaactactggcggcagctgctgaacgccaagctgattacccagagaaagttcgacaatctgaccaaggccgagagaggcggcctgagcgaactggataaggccggcttcatcaagagacagctggtggaaacccggcagatcacaaagcacgtggcacagatcctggactcccggatgaacactaagtacgacgagaatgacaagctgatccgggaagtgaaagtgatcaccctgaagtccaagctggtgtccgatttccggaaggatttccagttttacaaagtgcgcgagatcaacaactaccaccacgcccacgacgcctacctgaacgccgtcgtgggaaccgccctgatcaaaaagtaccctaagctggaaagcgagttcgtgtacggcgactacaaggtgtacgacgtgcggaagatgatcgccaagagcgagcaggaaatcggcaaggctaccgccaagtacttcttctacagcaacatcatgaactttttcaagaccgagattaccctggccaacggcgagatccggaagcggcctctgatcgagacaaacggcgaaaccggggagatcgtgtgggataagggccgggattttgccaccgtgcggaaagtgctgagcatgccccaagtgaatatcgtgaaaaagaccgaggtgcagacaggcggcttcagcaaagagtctatcctgcccaagaggaacagcgataagctgatcgccagaaagaaggactgggaccctaagaagtacggcggcttcgacagccccaccgtggcctattctgtgctggtggtggccaaagtggaaaagggcaagtccaagaaactgaagagtgtgaaagagctgctggggatcaccatcatggaaagaagcagcttcgagaagaatcccatcgactttctggaagccaagggctacaaagaagtgaaaaaggacctgatcatcaagctgcctaagtactccctgttcgagctggaaaacggccggaagagaatgctggcctctgccggcgaactgcagaagggaaacgaactggccctgccctccaaatatgtgaacttcctgtacctggccagccactatgagaagctgaagggctcccccgaggataatgagcagaaacagctgtttgtggaacagcacaagcactacctggacgagatcatcgagcagatcagcgagttctccaagagagtgatcctggccgacgctaatctggacaaagtgctgtccgcctacaacaagcaccgggataagcccatcagagagcaggccgagaatatcatccacctgtttaccctgaccaatctgggagcccctgccgccttcaagtactttgacaccaccatcgaccggaagaggtacaccagcaccaaagaggtgctggacgccaccctgatccaccagagcatcaccggcctgtacgagacacggatcgacctgtctcagctgggaggcgacagcggcgggagcggcgggagcggggggagcactaatctgagcgacatcattgagaaggagactgggaaacagctggtcattcaggagtccatcctgatgctgcctgaggaggtggaggaagtgatcggcaacaagccagagtctgacatcctggtgcacaccgcctacgacgagtccacagatgagaatgtgatgctgctgacctctgacgcccccgagtataagccttgggccctggtcatccaggattctaacggcgagaataagatcaagatgctgagcggaggatctggaggatctggaggcagcaccaacctgtctgacatcatcgagaaggagacaggcaagcagctggtcatccaggagagcatcctgatgctgcccgaagaagtcgaagaagtgatcggaaacaagcctgagagcggtatcctggtccataccgcctacgacgagagtaccgacgaaaatgtgatgctgctgacatccgacgccccagagtataagccctgggctctggtcatccaggattccaacggagagaacaaaatcaaaatgctgtctggcggctcaaaaagaaccgccgacggcagcgaatttgagaagcgacctgccgccacaaagaaggctggacaggctaagaagaagaaagattacaaagacgatgacgataagggatccggcgcaacaaacttctctctgctgaaacaagccggagatgtcgaagagaatcctggaccgatggccaagcctttgtctcaagaagaatccaccctcattgaaagagcaacggctacaatcaacagcatccccatctctgaagactacagcgtcgccagcgcagctctctctagcgacggccgcatcttcactggtgtcaatgtatatcattttactgggggaccttgtgcagaactcgtggtgctgggcactgctgctgctgcggcagctggcaacctgacttgtatcgtcgcgatcggaaatgagaacaggggcatcttgagcccctgcggacggtgccgacaggtgcttctcgatctgcatcctgggatcaaagccatagtgaaggacagtgatggacagccgacggcagttgggattcgtgaattgctgccctctggttatgtgtgggagggctaa

**g**, pLenti-xCas9-evoA-BE4max-Blast

gggcagagcgcacatcgcccacagtccccgagaagttggggggaggggtcggcaattgatccggtgcctagagaaggtggcgcggggtaaactgggaaagtgatgtcgtgtactggctccgcctttttcccgagggtgggggagaaccgtatataagtgcagtagtcgccgtgaacgttctttttcgcaacgggtttgccgccagaacacaggaccggttctagaatgaaacggacagccgacggaagcgagttcgagtcaccaaagaagaagcggaaagtcagttcaaagactgggcctgtcgccgtcgatccaaccctgcgccgccggattgaacctcacgagtttgaagtgttctttgacccccgggagctgagaaaggagacatgcctgctgtacgagatcaactggggaggcaggcactccatctggaggcacacctctcagaacacaaataagcacgtggaggtgaacttcatcgagaagtttaccacagagcggtacttctgccccaataccagatgtagcatcacatggtttctgagctggtccccttgcggagagtgtagcagggccatcaccgagttcctgtccagatacccaaatgtgacactgtttatctacatcgccaggctgtatcacctggcaaacccaaggaataggcagggcctgcgcgatctgatcagctccggcgtgaccatccagatcatgacagagcaggagtccggctactgctggcacaacttcgtgaattattctcctagcaacgagtcccactggcctaggtacccacacctgtgggtgcgcctgtacgtgctggagctgtattgcatcatcctgggcctgcccccttgtctgaatatcctgcggagaaagcagagccagctgacctcctttacaatcgccctgcagtcttgtcactatcagaggctgccaccccacatcctgtgggccacaggcctgaagtctggcggatctagcggaggatcttctggcagcgagacaccaggaacaagcgagtcagcaacaccagagagcagtggcggcagcagcggcggcagcgacaagaagtacagcatcggcctggCcatcggcaccaactctgtgggctgggccgtgatcaccgacgagtacaaggtgcccagcaagaaattcaaggtgctgggcaacaccgaccggcacagcatcaagaagaacctgatcggagccctgctgttcgacagcggcgaaacagccgaggccacccggctgaagagaaccgccagaagaagatacaccagacggaagaaccggatctgctatctgcaagagatcttcagcaacgagatggccaaggtggacgacagcttcttccacagactggaagagtccttcctggtggaagaggataagaagcacgagcggcaccccatcttcggcaacatcgtggacgaggtggcctaccacgagaagtaccccaccatctaccacctgagaaagaaactggtggacagcaccgacaaggccgacctgcggctgatctatctggccctggcccacatgatcaagttccggggccacttcctgatcgagggcgacctgaaccccgacaacagcgacgtggacaagctgttcatccagctggtgcagacctacaaccagctgttcgaggaaaaccccatcaacgccagcggcgtggacgccaaggccatcctgtctgccagactgagcaagagcagacggctggaaaatctgatcgcccagctgcccggcgagaagaagaatggcctgttcggaaacctgattgccctgagcctgggcctgacccccaacttcaagagcaacttcgacctggccgaggatAccaaactgcagctgagcaaggacacctacgacgacgacctggacaacctgctggcccagatcggcgaccagtacgccgacctgtttctggccgccaagaacctgtccgacgccatcctgctgagcgacatcctgagagtgaacaccgagatcaccaaggcccccctgagcgcctctatgatcaagCTatacgacgagcaccaccaggacctgaccctgctgaaagctctcgtgcggcagcagctgcctgagaagtacaaagagattttcttcgaccagagcaagaacggctacgccggctacattgacggcggagccagccaggaagagttctacaagttcatcaagcccatcctggaaaagatggacggcaccgaggaactgctcgtgaagctgaacagagaggacctgctgcggaagcagcggaccttcgacaacggcaTcatcccccaccagatccacctgggagagctgcacgccattctgcggcggcaggaagatttttacccattcctgaaggacaaccgggaaaagatcgagaagatcctgaccttccgcatcccctactacgtgggccctctggccaggggaaacagcagattcgcctggatgaccagaaagagcgaggaaaccatcaccccctggaacttcgagAaagtggtggacaagggcgcttccgcccagagcttcatcgagcggatgaccaacttcgataagaacctgcccaacgagaaggtgctgcccaagcacagcctgctgtacgagtacttcaccgtgtataacgagctgaccaaagtgaaatacgtgaccgagggaatgagaaagcccgccttcctgagcggcgaTcagaaaaaggccatcgtggacctgctgttcaagaccaaccggaaagtgaccgtgaagcagctgaaagaggactacttcaagaaaatcgagtgcttcgactccgtggaaatctccggcgtggaagatcggttcaacgcctccctgggcacataccacgatctgctgaaaattatcaaggacaaggacttcctggacaatgaggaaaacgaggacattctggaagatatcgtgctgaccctgacactgtttgaggacagagagatgatcgaggaacggctgaaaacctatgcccacctgttcgacgacaaagtgatgaagcagctgaagcggcggagatacaccggctggggcaggctgagccggaagctgatcaacggcatccgggacaagcagtccggcaagacaatcctggatttcctgaagtccgacggcttcgccaacagaaacttcatTcagctgatccacgacgacagcctgacctttaaagaggacatccagaaagcccaggtgtccggccagggcgatagcctgcacgagcacattgccaatctggccggcagccccgccattaagaagggcatcctgcagacagtgaaggtggtggacgagctcgtgaaagtgatgggccggcacaagcccgagaacatcgtgatcgaaatggccagagagaaccagaccacccagaagggacagaagaacagccgcgagagaatgaagcggatcgaagagggcatcaaagagctgggcagccagatcctgaaagaacaccccgtggaaaacacccagctgcagaacgagaagctgtacctgtactacctgcagaatgggcgggatatgtacgtggaccaggaactggacatcaaccggctgtccgactacgatgtggaccatatcgtgcctcagagctttctgaaggacgactccatcgacaacaaggtgctgaccagaagcgacaagaaccggggcaagagcgacaacgtgccctccgaagaggtcgtgaagaagatgaagaactactggcggcagctgctgaacgccaagctgattacccagagaaagttcgacaatctgaccaaggccgagagaggcggcctgagcgaactggataaggccggcttcatcaagagacagctggtggaaacccggcagatcacaaagcacgtggcacagatcctggactcccggatgaacactaagtacgacgagaatgacaagctgatccgggaagtgaaagtgatcaccctgaagtccaagctggtgtccgatttccggaaggatttccagttttacaaagtgcgcgagatcaacaactaccaccacgcccacgacgcctacctgaacgccgtcgtgggaaccgccctgatcaaaaagtaccctaagctggaaagcgagttcgtgtacggcgactacaaggtgtacgacgtgcggaagatgatcgccaagagcgagcaggaaatcggcaaggctaccgccaagtacttcttctacagcaacatcatgaactttttcaagaccgagattaccctggccaacggcgagatccggaagcggcctctgatcgagacaaacggcgaaaccggggagatcgtgtgggataagggccgggattttgccaccgtgcggaaagtgctgagcatgccccaagtgaatatcgtgaaaaagaccgaggtgcagacaggcggcttcagcaaagagtctatcctgcccaagaggaacagcgataagctgatcgccagaaagaaggactgggaccctaagaagtacggcggcttcgacagccccaccgtggcctattctgtgctggtggtggccaaagtggaaaagggcaagtccaagaaactgaagagtgtgaaagagctgctggggatcaccatcatggaaagaagcagcttcgagaagaatcccatcgactttctggaagccaagggctacaaagaagtgaaaaaggacctgatcatcaagctgcctaagtactccctgttcgagctggaaaacggccggaagagaatgctggcctctgccggcgTactgcagaagggaaacgaactggccctgccctccaaatatgtgaacttcctgtacctggccagccactatgagaagctgaagggctcccccgaggataatgagcagaaacagctgtttgtggaacagcacaagcactacctggacgagatcatcgagcagatcagcgagttctccaagagagtgatcctggccgacgctaatctggacaaagtgctgtccgcctacaacaagcaccgggataagcccatcagagagcaggccgagaatatcatccacctgtttaccctgaccaatctgggagcccctgccgccttcaagtactttgacaccaccatcgaccggaagaggtacaccagcaccaaagaggtgctggacgccaccctgatccaccagagcatcaccggcctgtacgagacacggatcgacctgtctcagctgggaggcgacagcggcgggagcggcgggagcggggggagcactaatctgagcgacatcattgagaaggagactgggaaacagctggtcattcaggagtccatcctgatgctgcctgaggaggtggaggaagtgatcggcaacaagccagagtctgacatcctggtgcacaccgcctacgacgagtccacagatgagaatgtgatgctgctgacctctgacgcccccgagtataagccttgggccctggtcatccaggattctaacggcgagaataagatcaagatgctgagcggaggatctggaggatctggaggcagcaccaacctgtctgacatcatcgagaaggagacaggcaagcagctggtcatccaggagagcatcctgatgctgcccgaagaagtcgaagaagtgatcggaaacaagcctgagagcggtatcctggtccataccgcctacgacgagagtaccgacgaaaatgtgatgctgctgacatccgacgccccagagtataagccctgggctctggtcatccaggattccaacggagagaacaaaatcaaaatgctgtctggcggctcaaaaagaaccgccgacggcagcgaatttgagaagcgacctgccgccacaaagaaggctggacaggctaagaagaagaaagattacaaagacgatgacgataagggatccggcgcaacaaacttctctctgctgaaacaagccggagatgtcgaagagaatcctggaccgatggccaagcctttgtctcaagaagaatccaccctcattgaaagagcaacggctacaatcaacagcatccccatctctgaagactacagcgtcgccagcgcagctctctctagcgacggccgcatcttcactggtgtcaatgtatatcattttactgggggaccttgtgcagaactcgtggtgctgggcactgctgctgctgcggcagctggcaacctgacttgtatcgtcgcgatcggaaatgagaacaggggcatcttgagcccctgcggacggtgccgacaggtgcttctcgatctgcatcctgggatcaaagccatagtgaaggacagtgatggacagccgacggcagttgggattcgtgaattgctgccctctggttatgtgtgggagggctaa

**h**, pLenti-Cas9-NG-evoA-BE4max-Blast

gggcagagcgcacatcgcccacagtccccgagaagttggggggaggggtcggcaattgatccggtgcctagagaaggtggcgcggggtaaactgggaaagtgatgtcgtgtactggctccgcctttttcccgagggtgggggagaaccgtatataagtgcagtagtcgccgtgaacgttctttttcgcaacgggtttgccgccagaacacaggaccggttctagaatgaaacggacagccgacggaagcgagttcgagtcaccaaagaagaagcggaaagtcagttcaaagactgggcctgtcgccgtcgatccaaccctgcgccgccggattgaacctcacgagtttgaagtgttctttgacccccgggagctgagaaaggagacatgcctgctgtacgagatcaactggggaggcaggcactccatctggaggcacacctctcagaacacaaataagcacgtggaggtgaacttcatcgagaagtttaccacagagcggtacttctgccccaataccagatgtagcatcacatggtttctgagctggtccccttgcggagagtgtagcagggccatcaccgagttcctgtccagatacccaaatgtgacactgtttatctacatcgccaggctgtatcacctggcaaacccaaggaataggcagggcctgcgcgatctgatcagctccggcgtgaccatccagatcatgacagagcaggagtccggctactgctggcacaacttcgtgaattattctcctagcaacgagtcccactggcctaggtacccacacctgtgggtgcgcctgtacgtgctggagctgtattgcatcatcctgggcctgcccccttgtctgaatatcctgcggagaaagcagagccagctgacctcctttacaatcgccctgcagtcttgtcactatcagaggctgccaccccacatcctgtgggccacaggcctgaagtctggcggatctagcggaggatcttctggcagcgagacaccaggaacaagcgagtcagcaacaccagagagcagtggcggcagcagcggcggcagcgacaagaagtacagcatcggcctggccatcggcaccaactctgtgggctgggccgtgatcaccgacgagtacaaggtgcccagcaagaaattcaaggtgctgggcaacaccgaccggcacagcatcaagaagaacctgatcggagccctgctgttcgacagcggcgaaacagccgaggccacccggctgaagagaaccgccagaagaagatacaccagacggaagaaccggatctgctatctgcaagagatcttcagcaacgagatggccaaggtggacgacagcttcttccacagactggaagagtccttcctggtggaagaggataagaagcacgagcggcaccccatcttcggcaacatcgtggacgaggtggcctaccacgagaagtaccccaccatctaccacctgagaaagaaactggtggacagcaccgacaaggccgacctgcggctgatctatctggccctggcccacatgatcaagttccggggccacttcctgatcgagggcgacctgaaccccgacaacagcgacgtggacaagctgttcatccagctggtgcagacctacaaccagctgttcgaggaaaaccccatcaacgccagcggcgtggacgccaaggccatcctgtctgccagactgagcaagagcagacggctggaaaatctgatcgcccagctgcccggcgagaagaagaatggcctgttcggaaacctgattgccctgagcctgggcctgacccccaacttcaagagcaacttcgacctggccgaggatgccaaactgcagctgagcaaggacacctacgacgacgacctggacaacctgctggcccagatcggcgaccagtacgccgacctgtttctggccgccaagaacctgtccgacgccatcctgctgagcgacatcctgagagtgaacaccgagatcaccaaggcccccctgagcgcctctatgatcaagagatacgacgagcaccaccaggacctgaccctgctgaaagctctcgtgcggcagcagctgcctgagaagtacaaagagattttcttcgaccagagcaagaacggctacgccggctacattgacggcggagccagccaggaagagttctacaagttcatcaagcccatcctggaaaagatggacggcaccgaggaactgctcgtgaagctgaacagagaggacctgctgcggaagcagcggaccttcgacaacggcagcatcccccaccagatccacctgggagagctgcacgccattctgcggcggcaggaagatttttacccattcctgaaggacaaccgggaaaagatcgagaagatcctgaccttccgcatcccctactacgtgggccctctggccaggggaaacagcagattcgcctggatgaccagaaagagcgaggaaaccatcaccccctggaacttcgaggaagtggtggacaagggcgcttccgcccagagcttcatcgagcggatgaccaacttcgataagaacctgcccaacgagaaggtgctgcccaagcacagcctgctgtacgagtacttcaccgtgtataacgagctgaccaaagtgaaatacgtgaccgagggaatgagaaagcccgccttcctgagcggcgagcagaaaaaggccatcgtggacctgctgttcaagaccaaccggaaagtgaccgtgaagcagctgaaagaggactacttcaagaaaatcgagtgcttcgactccgtggaaatctccggcgtggaagatcggttcaacgcctccctgggcacataccacgatctgctgaaaattatcaaggacaaggacttcctggacaatgaggaaaacgaggacattctggaagatatcgtgctgaccctgacactgtttgaggacagagagatgatcgaggaacggctgaaaacctatgcccacctgttcgacgacaaagtgatgaagcagctgaagcggcggagatacaccggctggggcaggctgagccggaagctgatcaacggcatccgggacaagcagtccggcaagacaatcctggatttcctgaagtccgacggcttcgccaacagaaacttcatgcagctgatccacgacgacagcctgacctttaaagaggacatccagaaagcccaggtgtccggccagggcgatagcctgcacgagcacattgccaatctggccggcagccccgccattaagaagggcatcctgcagacagtgaaggtggtggacgagctcgtgaaagtgatgggccggcacaagcccgagaacatcgtgatcgaaatggccagagagaaccagaccacccagaagggacagaagaacagccgcgagagaatgaagcggatcgaagagggcatcaaagagctgggcagccagatcctgaaagaacaccccgtggaaaacacccagctgcagaacgagaagctgtacctgtactacctgcagaatgggcgggatatgtacgtggaccaggaactggacatcaaccggctgtccgactacgatgtggaccatatcgtgcctcagagctttctgaaggacgactccatcgacaacaaggtgctgaccagaagcgacaagaaccggggcaagagcgacaacgtgccctccgaagaggtcgtgaagaagatgaagaactactggcggcagctgctgaacgccaagctgattacccagagaaagttcgacaatctgaccaaggccgagagaggcggcctgagcgaactggataaggccggcttcatcaagagacagctggtggaaacccggcagatcacaaagcacgtggcacagatcctggactcccggatgaacactaagtacgacgagaatgacaagctgatccgggaagtgaaagtgatcaccctgaagtccaagctggtgtccgatttccggaaggatttccagttttacaaagtgcgcgagatcaacaactaccaccacgcccacgacgcctacctgaacgccgtcgtgggaaccgccctgatcaaaaagtaccctaagctggaaagcgagttcgtgtacggcgactacaaggtgtacgacgtgcggaagatgatcgccaagagcgagcaggaaatcggcaaggctaccgccaagtacttcttctacagcaacatcatgaactttttcaagaccgagattaccctggccaacggcgagatccggaagcggcctctgatcgagacaaacggcgaaaccggggagatcgtgtgggataagggccgggattttgccaccgtgcggaaagtgctgagcatgccccaagtgaatatcgtgaaaaagaccgaggtgcagacaggcggcttcagcaaagagtctatccGgcccaagaggaacagcgataagctgatcgccagaaagaaggactgggaccctaagaagtacggcggcttcgTcagccccaccgtggcctattctgtgctggtggtggccaaagtggaaaagggcaagtccaagaaactgaagagtgtgaaagagctgctggggatcaccatcatggaaagaagcagcttcgagaagaatcccatcgactttctggaagccaagggctacaaagaagtgaaaaaggacctgatcatcaagctgcctaagtactccctgttcgagctggaaaacggccggaagagaatgctggcctctgccCgcTTCctgcagaagggaaacgaactggccctgccctccaaatatgtgaacttcctgtacctggccagccactatgagaagctgaagggctcccccgaggataatgagcagaaacagctgtttgtggaacagcacaagcactacctggacgagatcatcgagcagatcagcgagttctccaagagagtgatcctggccgacgctaatctggacaaagtgctgtccgcctacaacaagcaccgggataagcccatcagagagcaggccgagaatatcatccacctgtttaccctgaccaatctgggagcccctCGcgccttcaagtactttgacaccaccatcgaccggaagGTgtacaGGagcaccaaagaggtgctggacgccaccctgatccaccagagcatcaccggcctgtacgagacacggatcgacctgtctcagctgggaggcgacagcggcgggagcggcgggagcggggggagcactaatctgagcgacatcattgagaaggagactgggaaacagctggtcattcaggagtccatcctgatgctgcctgaggaggtggaggaagtgatcggcaacaagccagagtctgacatcctggtgcacaccgcctacgacgagtccacagatgagaatgtgatgctgctgacctctgacgcccccgagtataagccttgggccctggtcatccaggattctaacggcgagaataagatcaagatgctgagcggaggatctggaggatctggaggcagcaccaacctgtctgacatcatcgagaaggagacaggcaagcagctggtcatccaggagagcatcctgatgctgcccgaagaagtcgaagaagtgatcggaaacaagcctgagagcggtatcctggtccataccgcctacgacgagagtaccgacgaaaatgtgatgctgctgacatccgacgccccagagtataagccctgggctctggtcatccaggattccaacggagagaacaaaatcaaaatgctgtctggcggctcaaaaagaaccgccgacggcagcgaatttgagaagcgacctgccgccacaaagaaggctggacaggctaagaagaagaaagattacaaagacgatgacgataagggatccggcgcaacaaacttctctctgctgaaacaagccggagatgtcgaagagaatcctggaccgatggccaagcctttgtctcaagaagaatccaccctcattgaaagagcaacggctacaatcaacagcatccccatctctgaagactacagcgtcgccagcgcagctctctctagcgacggccgcatcttcactggtgtcaatgtatatcattttactgggggaccttgtgcagaactcgtggtgctgggcactgctgctgctgcggcagctggcaacctgacttgtatcgtcgcgatcggaaatgagaacaggggcatcttgagcccctgcggacggtgccgacaggtgcttctcgatctgcatcctgggatcaaagccatagtgaaggacagtgatggacagccgacggcagttgggattcgtgaattgctgccctctggttatgtgtgggagggctaa

**i**, pLenti-SpG-evoA-BE4max-Blast

gggcagagcgcacatcgcccacagtccccgagaagttggggggaggggtcggcaattgatccggtgcctagagaaggtggcgcggggtaaactgggaaagtgatgtcgtgtactggctccgcctttttcccgagggtgggggagaaccgtatataagtgcagtagtcgccgtgaacgttctttttcgcaacgggtttgccgccagaacacaggaccggttctagaatgaaacggacagccgacggaagcgagttcgagtcaccaaagaagaagcggaaagtcagttcaaagactgggcctgtcgccgtcgatccaaccctgcgccgccggattgaacctcacgagtttgaagtgttctttgacccccgggagctgagaaaggagacatgcctgctgtacgagatcaactggggaggcaggcactccatctggaggcacacctctcagaacacaaataagcacgtggaggtgaacttcatcgagaagtttaccacagagcggtacttctgccccaataccagatgtagcatcacatggtttctgagctggtccccttgcggagagtgtagcagggccatcaccgagttcctgtccagatacccaaatgtgacactgtttatctacatcgccaggctgtatcacctggcaaacccaaggaataggcagggcctgcgcgatctgatcagctccggcgtgaccatccagatcatgacagagcaggagtccggctactgctggcacaacttcgtgaattattctcctagcaacgagtcccactggcctaggtacccacacctgtgggtgcgcctgtacgtgctggagctgtattgcatcatcctgggcctgcccccttgtctgaatatcctgcggagaaagcagagccagctgacctcctttacaatcgccctgcagtcttgtcactatcagaggctgccaccccacatcctgtgggccacaggcctgaagtctggcggatctagcggaggatcttctggcagcgagacaccaggaacaagcgagtcagcaacaccagagagcagtggcggcagcagcggcggcagcgacaagaagtacagcatcggcctggccatcggcaccaactctgtgggctgggccgtgatcaccgacgagtacaaggtgcccagcaagaaattcaaggtgctgggcaacaccgaccggcacagcatcaagaagaacctgatcggagccctgctgttcgacagcggcgaaacagccgaggccacccggctgaagagaaccgccagaagaagatacaccagacggaagaaccggatctgctatctgcaagagatcttcagcaacgagatggccaaggtggacgacagcttcttccacagactggaagagtccttcctggtggaagaggataagaagcacgagcggcaccccatcttcggcaacatcgtggacgaggtggcctaccacgagaagtaccccaccatctaccacctgagaaagaaactggtggacagcaccgacaaggccgacctgcggctgatctatctggccctggcccacatgatcaagttccggggccacttcctgatcgagggcgacctgaaccccgacaacagcgacgtggacaagctgttcatccagctggtgcagacctacaaccagctgttcgaggaaaaccccatcaacgccagcggcgtggacgccaaggccatcctgtctgccagactgagcaagagcagacggctggaaaatctgatcgcccagctgcccggcgagaagaagaatggcctgttcggaaacctgattgccctgagcctgggcctgacccccaacttcaagagcaacttcgacctggccgaggatgccaaactgcagctgagcaaggacacctacgacgacgacctggacaacctgctggcccagatcggcgaccagtacgccgacctgtttctggccgccaagaacctgtccgacgccatcctgctgagcgacatcctgagagtgaacaccgagatcaccaaggcccccctgagcgcctctatgatcaagagatacgacgagcaccaccaggacctgaccctgctgaaagctctcgtgcggcagcagctgcctgagaagtacaaagagattttcttcgaccagagcaagaacggctacgccggctacattgacggcggagccagccaggaagagttctacaagttcatcaagcccatcctggaaaagatggacggcaccgaggaactgctcgtgaagctgaacagagaggacctgctgcggaagcagcggaccttcgacaacggcagcatcccccaccagatccacctgggagagctgcacgccattctgcggcggcaggaagatttttacccattcctgaaggacaaccgggaaaagatcgagaagatcctgaccttccgcatcccctactacgtgggccctctggccaggggaaacagcagattcgcctggatgaccagaaagagcgaggaaaccatcaccccctggaacttcgaggaagtggtggacaagggcgcttccgcccagagcttcatcgagcggatgaccaacttcgataagaacctgcccaacgagaaggtgctgcccaagcacagcctgctgtacgagtacttcaccgtgtataacgagctgaccaaagtgaaatacgtgaccgagggaatgagaaagcccgccttcctgagcggcgagcagaaaaaggccatcgtggacctgctgttcaagaccaaccggaaagtgaccgtgaagcagctgaaagaggactacttcaagaaaatcgagtgcttcgactccgtggaaatctccggcgtggaagatcggttcaacgcctccctgggcacataccacgatctgctgaaaattatcaaggacaaggacttcctggacaatgaggaaaacgaggacattctggaagatatcgtgctgaccctgacactgtttgaggacagagagatgatcgaggaacggctgaaaacctatgcccacctgttcgacgacaaagtgatgaagcagctgaagcggcggagatacaccggctggggcaggctgagccggaagctgatcaacggcatccgggacaagcagtccggcaagacaatcctggatttcctgaagtccgacggcttcgccaacagaaacttcatgcagctgatccacgacgacagcctgacctttaaagaggacatccagaaagcccaggtgtccggccagggcgatagcctgcacgagcacattgccaatctggccggcagccccgccattaagaagggcatcctgcagacagtgaaggtggtggacgagctcgtgaaagtgatgggccggcacaagcccgagaacatcgtgatcgaaatggccagagagaaccagaccacccagaagggacagaagaacagccgcgagagaatgaagcggatcgaagagggcatcaaagagctgggcagccagatcctgaaagaacaccccgtggaaaacacccagctgcagaacgagaagctgtacctgtactacctgcagaatgggcgggatatgtacgtggaccaggaactggacatcaaccggctgtccgactacgatgtggaccatatcgtgcctcagagctttctgaaggacgactccatcgacaacaaggtgctgaccagaagcgacaagaaccggggcaagagcgacaacgtgccctccgaagaggtcgtgaagaagatgaagaactactggcggcagctgctgaacgccaagctgattacccagagaaagttcgacaatctgaccaaggccgagagaggcggcctgagcgaactggataaggccggcttcatcaagagacagctggtggaaacccggcagatcacaaagcacgtggcacagatcctggactcccggatgaacactaagtacgacgagaatgacaagctgatccgggaagtgaaagtgatcaccctgaagtccaagctggtgtccgatttccggaaggatttccagttttacaaagtgcgcgagatcaacaactaccaccacgcccacgacgcctacctgaacgccgtcgtgggaaccgccctgatcaaaaagtaccctaagctggaaagcgagttcgtgtacggcgactacaaggtgtacgacgtgcggaagatgatcgccaagagcgagcaggaaatcggcaaggctaccgccaagtacttcttctacagcaacatcatgaactttttcaagaccgagattaccctggccaacggcgagatccggaagcggcctctgatcgagacaaacggcgaaaccggggagatcgtgtgggataagggccgggattttgccaccgtgcggaaagtgctgagcatgccccaagtgaatatcgtgaaaaagaccgaggtgcagacaggcggcttcagcaaagagtctatcctgcccaagaggaacagcgataagctgatcgccagaaagaaggactgggaccctaagaagtacggcggcttcCTGTgGcccaccgtggcctattctgtgctggtggtggccaaagtggaaaagggcaagtccaagaaactgaagagtgtgaaagagctgctggggatcaccatcatggaaagaagcagcttcgagaagaatcccatcgactttctggaagccaagggctacaaagaagtgaaaaaggacctgatcatcaagctgcctaagtactccctgttcgagctggaaaacggccggaagagaatgctggcctctgccAAGCaGctgcagaagggaaacgaactggccctgccctccaaatatgtgaacttcctgtacctggccagccactatgagaagctgaagggctcccccgaggataatgagcagaaacagctgtttgtggaacagcacaagcactacctggacgagatcatcgagcagatcagcgagttctccaagagagtgatcctggccgacgctaatctggacaaagtgctgtccgcctacaacaagcaccgggataagcccatcagagagcaggccgagaatatcatccacctgtttaccctgaccaatctgggagcccctgccgccttcaagtactttgacaccaccatcgaccggaagCAgtacaGAagcaccaaagaggtgctggacgccaccctgatccaccagagcatcaccggcctgtacgagacacggatcgacctgtctcagctgggaggcgacagcggcgggagcggcgggagcggggggagcactaatctgagcgacatcattgagaaggagactgggaaacagctggtcattcaggagtccatcctgatgctgcctgaggaggtggaggaagtgatcggcaacaagccagagtctgacatcctggtgcacaccgcctacgacgagtccacagatgagaatgtgatgctgctgacctctgacgcccccgagtataagccttgggccctggtcatccaggattctaacggcgagaataagatcaagatgctgagcggaggatctggaggatctggaggcagcaccaacctgtctgacatcatcgagaaggagacaggcaagcagctggtcatccaggagagcatcctgatgctgcccgaagaagtcgaagaagtgatcggaaacaagcctgagagcggtatcctggtccataccgcctacgacgagagtaccgacgaaaatgtgatgctgctgacatccgacgccccagagtataagccctgggctctggtcatccaggattccaacggagagaacaaaatcaaaatgctgtctggcggctcaaaaagaaccgccgacggcagcgaatttgagaagcgacctgccgccacaaagaaggctggacaggctaagaagaagaaagattacaaagacgatgacgataagggatccggcgcaacaaacttctctctgctgaaacaagccggagatgtcgaagagaatcctggaccgatggccaagcctttgtctcaagaagaatccaccctcattgaaagagcaacggctacaatcaacagcatccccatctctgaagactacagcgtcgccagcgcagctctctctagcgacggccgcatcttcactggtgtcaatgtatatcattttactgggggaccttgtgcagaactcgtggtgctgggcactgctgctgctgcggcagctggcaacctgacttgtatcgtcgcgatcggaaatgagaacaggggcatcttgagcccctgcggacggtgccgacaggtgcttctcgatctgcatcctgggatcaaagccatagtgaaggacagtgatggacagccgacggcagttgggattcgtgaattgctgccctctggttatgtgtgggagggctaa

**j**, pLenti-SpRY-evoA-BE4max-Blast

gggcagagcgcacatcgcccacagtccccgagaagttggggggaggggtcggcaattgatccggtgcctagagaaggtggcgcggggtaaactgggaaagtgatgtcgtgtactggctccgcctttttcccgagggtgggggagaaccgtatataagtgcagtagtcgccgtgaacgttctttttcgcaacgggtttgccgccagaacacaggaccggttctagaatgaaacggacagccgacggaagcgagttcgagtcaccaaagaagaagcggaaagtcagttcaaagactgggcctgtcgccgtcgatccaaccctgcgccgccggattgaacctcacgagtttgaagtgttctttgacccccgggagctgagaaaggagacatgcctgctgtacgagatcaactggggaggcaggcactccatctggaggcacacctctcagaacacaaataagcacgtggaggtgaacttcatcgagaagtttaccacagagcggtacttctgccccaataccagatgtagcatcacatggtttctgagctggtccccttgcggagagtgtagcagggccatcaccgagttcctgtccagatacccaaatgtgacactgtttatctacatcgccaggctgtatcacctggcaaacccaaggaataggcagggcctgcgcgatctgatcagctccggcgtgaccatccagatcatgacagagcaggagtccggctactgctggcacaacttcgtgaattattctcctagcaacgagtcccactggcctaggtacccacacctgtgggtgcgcctgtacgtgctggagctgtattgcatcatcctgggcctgcccccttgtctgaatatcctgcggagaaagcagagccagctgacctcctttacaatcgccctgcagtcttgtcactatcagaggctgccaccccacatcctgtgggccacaggcctgaagtctggcggatctagcggaggatcttctggcagcgagacaccaggaacaagcgagtcagcaacaccagagagcagtggcggcagcagcggcggcagcgacaagaagtacagcatcggcctggccatcggcaccaactctgtgggctgggccgtgatcaccgacgagtacaaggtgcccagcaagaaattcaaggtgctgggcaacaccgaccggcacagcatcaagaagaacctgatcggagccctgctgttcgacagcggcgaaacagccgagAGAacccggctgaagagaaccgccagaagaagatacaccagacggaagaaccggatctgctatctgcaagagatcttcagcaacgagatggccaaggtggacgacagcttcttccacagactggaagagtccttcctggtggaagaggataagaagcacgagcggcaccccatcttcggcaacatcgtggacgaggtggcctaccacgagaagtaccccaccatctaccacctgagaaagaaactggtggacagcaccgacaaggccgacctgcggctgatctatctggccctggcccacatgatcaagttccggggccacttcctgatcgagggcgacctgaaccccgacaacagcgacgtggacaagctgttcatccagctggtgcagacctacaaccagctgttcgaggaaaaccccatcaacgccagcggcgtggacgccaaggccatcctgtctgccagactgagcaagagcagacggctggaaaatctgatcgcccagctgcccggcgagaagaagaatggcctgttcggaaacctgattgccctgagcctgggcctgacccccaacttcaagagcaacttcgacctggccgaggatgccaaactgcagctgagcaaggacacctacgacgacgacctggacaacctgctggcccagatcggcgaccagtacgccgacctgtttctggccgccaagaacctgtccgacgccatcctgctgagcgacatcctgagagtgaacaccgagatcaccaaggcccccctgagcgcctctatgatcaagagatacgacgagcaccaccaggacctgaccctgctgaaagctctcgtgcggcagcagctgcctgagaagtacaaagagattttcttcgaccagagcaagaacggctacgccggctacattgacggcggagccagccaggaagagttctacaagttcatcaagcccatcctggaaaagatggacggcaccgaggaactgctcgtgaagctgaacagagaggacctgctgcggaagcagcggaccttcgacaacggcagcatcccccaccagatccacctgggagagctgcacgccattctgcggcggcaggaagatttttacccattcctgaaggacaaccgggaaaagatcgagaagatcctgaccttccgcatcccctactacgtgggccctctggccaggggaaacagcagattcgcctggatgaccagaaagagcgaggaaaccatcaccccctggaacttcgaggaagtggtggacaagggcgcttccgcccagagcttcatcgagcggatgaccaacttcgataagaacctgcccaacgagaaggtgctgcccaagcacagcctgctgtacgagtacttcaccgtgtataacgagctgaccaaagtgaaatacgtgaccgagggaatgagaaagcccgccttcctgagcggcgagcagaaaaaggccatcgtggacctgctgttcaagaccaaccggaaagtgaccgtgaagcagctgaaagaggactacttcaagaaaatcgagtgcttcgactccgtggaaatctccggcgtggaagatcggttcaacgcctccctgggcacataccacgatctgctgaaaattatcaaggacaaggacttcctggacaatgaggaaaacgaggacattctggaagatatcgtgctgaccctgacactgtttgaggacagagagatgatcgaggaacggctgaaaacctatgcccacctgttcgacgacaaagtgatgaagcagctgaagcggcggagatacaccggctggggcaggctgagccggaagctgatcaacggcatccgggacaagcagtccggcaagacaatcctggatttcctgaagtccgacggcttcgccaacagaaacttcatgcagctgatccacgacgacagcctgacctttaaagaggacatccagaaagcccaggtgtccggccagggcgatagcctgcacgagcacattgccaatctggccggcagccccgccattaagaagggcatcctgcagacagtgaaggtggtggacgagctcgtgaaagtgatgggccggcacaagcccgagaacatcgtgatcgaaatggccagagagaaccagaccacccagaagggacagaagaacagccgcgagagaatgaagcggatcgaagagggcatcaaagagctgggcagccagatcctgaaagaacaccccgtggaaaacacccagctgcagaacgagaagctgtacctgtactacctgcagaatgggcgggatatgtacgtggaccaggaactggacatcaaccggctgtccgactacgatgtggaccatatcgtgcctcagagctttctgaaggacgactccatcgacaacaaggtgctgaccagaagcgacaagaaccggggcaagagcgacaacgtgccctccgaagaggtcgtgaagaagatgaagaactactggcggcagctgctgaacgccaagctgattacccagagaaagttcgacaatctgaccaaggccgagagaggcggcctgagcgaactggataaggccggcttcatcaagagacagctggtggaaacccggcagatcacaaagcacgtggcacagatcctggactcccggatgaacactaagtacgacgagaatgacaagctgatccgggaagtgaaagtgatcaccctgaagtccaagctggtgtccgatttccggaaggatttccagttttacaaagtgcgcgagatcaacaactaccaccacgcccacgacgcctacctgaacgccgtcgtgggaaccgccctgatcaaaaagtaccctaagctggaaagcgagttcgtgtacggcgactacaaggtgtacgacgtgcggaagatgatcgccaagagcgagcaggaaatcggcaaggctaccgccaagtacttcttctacagcaacatcatgaactttttcaagaccgagattaccctggccaacggcgagatccggaagcggcctctgatcgagacaaacggcgaaaccggggagatcgtgtgggataagggccgggattttgccaccgtgcggaaagtgctgagcatgccccaagtgaatatcgtgaaaaagaccgaggtgcagacaggcggcttcagcaaagagtctatcAGAcccaagaggaacagcgataagctgatcgccagaaagaaggactgggaccctaagaagtacggcggcttcCTGTgGcccaccgtggcctattctgtgctggtggtggccaaagtggaaaagggcaagtccaagaaactgaagagtgtgaaagagctgctggggatcaccatcatggaaagaagcagcttcgagaagaatcccatcgactttctggaagccaagggctacaaagaagtgaaaaaggacctgatcatcaagctgcctaagtactccctgttcgagctggaaaacggccggaagagaatgctggcctctgccAAGCaGctgcagaagggaaacgaactggccctgccctccaaatatgtgaacttcctgtacctggccagccactatgagaagctgaagggctcccccgaggataatgagcagaaacagctgtttgtggaacagcacaagcactacctggacgagatcatcgagcagatcagcgagttctccaagagagtgatcctggccgacgctaatctggacaaagtgctgtccgcctacaacaagcaccgggataagcccatcagagagcaggccgagaatatcatccacctgtttaccctgaccaGActgggagcccctAGAgccttcaagtactttgacaccaccatcgaccCCaagCAgtacaGAagcaccaaagaggtgctggacgccaccctgatccaccagagcatcaccggcctgtacgagacacggatcgacctgtctcagctgggaggcgacagcggcgggagcggcgggagcggggggagcactaatctgagcgacatcattgagaaggagactgggaaacagctggtcattcaggagtccatcctgatgctgcctgaggaggtggaggaagtgatcggcaacaagccagagtctgacatcctggtgcacaccgcctacgacgagtccacagatgagaatgtgatgctgctgacctctgacgcccccgagtataagccttgggccctggtcatccaggattctaacggcgagaataagatcaagatgctgagcggaggatctggaggatctggaggcagcaccaacctgtctgacatcatcgagaaggagacaggcaagcagctggtcatccaggagagcatcctgatgctgcccgaagaagtcgaagaagtgatcggaaacaagcctgagagcggtatcctggtccataccgcctacgacgagagtaccgacgaaaatgtgatgctgctgacatccgacgccccagagtataagccctgggctctggtcatccaggattccaacggagagaacaaaatcaaaatgctgtctggcggctcaaaaagaaccgccgacggcagcgaatttgagaagcgacctgccgccacaaagaaggctggacaggctaagaagaagaaagattacaaagacgatgacgataagggatccggcgcaacaaacttctctctgctgaaacaagccggagatgtcgaagagaatcctggaccgatggccaagcctttgtctcaagaagaatccaccctcattgaaagagcaacggctacaatcaacagcatccccatctctgaagactacagcgtcgccagcgcagctctctctagcgacggccgcatcttcactggtgtcaatgtatatcattttactgggggaccttgtgcagaactcgtggtgctgggcactgctgctgctgcggcagctggcaacctgacttgtatcgtcgcgatcggaaatgagaacaggggcatcttgagcccctgcggacggtgccgacaggtgcttctcgatctgcatcctgggatcaaagccatagtgaaggacagtgatggacagccgacggcagttgggattcgtgaattgctgccctctggttatgtgtgggagggctaa
